# A microbiota membrane disrupter disseminates to the pancreas and increases β-cell mass

**DOI:** 10.1101/2022.03.24.485696

**Authors:** Jennifer Hampton Hill, Michelle Sconce Massaquoi, Emily Goers Sweeney, Elena S. Wall, Philip Jahl, Rickesha Bell, Karen Kallio, Daniel Derrick, L. Charles Murtaugh, Raghuveer Parthasarathy, S. James Remington, June L. Round, Karen Guillemin

**Author notes:** These authors contributed equally to this work (Michelle Sconce Massaquoi, Emily Goers Sweeney).

## Abstract

Microbiome dysbiosis is a feature of diabetes, but how microbial products influence insulin production is poorly understood. Here we report the mechanism of BefA, a microbiome-derived protein that increases proliferation of insulin-producing β-cells during pancreatic development in gnotobiotic zebrafish and mice. BefA disseminates systemically via multiple anatomic routes to act directly on pancreatic islets. We report the structure of BefA, containing a lipid-binding SYLF domain, and demonstrate that it permeabilizes synthetic liposomes and bacterial membranes. A BefA mutant impaired in membrane disruption fails to expand β-cells whereas the pore-forming host defense protein, Reg3, stimulates β-cell proliferation. Our work demonstrates that membrane permeabilization by microbiome-derived and host defense proteins is necessary and sufficient for β-cell expansion during pancreas development, thereby connecting microbiome composition with diabetes risk.

## Introduction

β-cells are an essential pancreatic endocrine cell type necessary for producing insulin to modulate blood glucose levels. Patients with diabetes lack functional β-cells due to either autoimmune destruction or long-term insulin resistance. Strategies for replacement of lost β-cells are sought as a means of treatment, particularly for type 1 diabetes (T1D), to restore endogenous insulin production (Aguayo-Mazzucato and Bonner-Weir, 2018). Extensive research has focused on understanding endogenous pathways of β-cell development and proliferation to harness these existing mechanisms for β-cell replacement therapies (Aguayo-Mazzucato and Bonner-Weir, 2018; Murtaugh, 2007). One example of an initially promising class of potential therapeutics was the **Reg**enerating islet-derived proteins, originally discovered in pancreatic tissue for their upregulation during the regenerative response of pancreatic β-cells (Terazono et al., 1988). These proteins however, were found to have pleiotropic effects, causing proliferation of several other tissues such as exocrine cells, intestinal epithelium, hepatocytes, and nerves [reviewed in (Chen et al., 2019)]. Subsequent to their characterization in the context of the pancreas, the Reg3 proteins were found to be potent pore-forming antimicrobial host defense proteins, essential for the maintenance of intestinal barrier function (Cash et al., 2006a; Mukherjee et al., 2014; Vaishnava et al., 2011). More recently, disease-protective microbiomes in the NOD mouse model of T1D were found to induce Reg3 expression (Zhang et al., 2021). However, no connections have been made between Reg3’s pore forming activity and its impact on the pancreas.

We recently showed that the resident microbiota promotes β-cell development in zebrafish larvae and discovered a secreted bacterial protein that could rescue germ free (GF) larval β-cell development, which we named BefA, for Beta-cell Expansion Factor A (Hill et al., 2016). BefA is a 29 kDa protein produced by a limited number of intestinal bacteria from various hosts including zebrafish, mice and humans (Hill et al., 2016). A BefA homolog from a human-derived *Klebsiella*, with only 34% amino acid sequence similarity to our original *Aeromonas veronii*-derived BefA, was sufficient to rescue GF zebrafish β-cell development (Hill et al., 2016). This result suggested that the mechanism by which BefA stimulates expansion of insulin-producing cells is mediated through structural features of the protein shared across distant homologs. It also raised the possibility that these features of BefA homologs from diverse host-associated bacteria may stimulate β-cell expansion across multiple host species, a question we explore in the current study.

Analysis of the amino acid sequence of the original BefA isolate and its functional homologs revealed little about the proteins’ activities, but uncovered sequence similarity to a lipid-binding SYLF domain (named for the first four proteins found to contain it, **S**H3YL1, **Y**sc84p, **L**sb4p, and **F**YVE), which is prevalent across the kingdoms of life (Hasegawa et al., 2011). Four eukaryotic SYLF-containing proteins, found in yeast, plants, and mammals, have been studied to date, and all are associated with cellular or organellar membranes (Dewar et al., 2002; Hasegawa et al., 2011; Robertson et al., 2009a; Sutipatanasomboon et al., 2017; Tonikian et al., 2009; Urbanek et al., 2015a; Wywial and Singh, 2010). Here we describe the atomic structure and biochemical features of a bacterial SYLF-domain protein. We find that BefA has an intrinsic affinity for membranes, inducing vesiculation and leakiness of synthetic membranes as well as loss of integrity of bacterial cells. We further identify a point mutation within the SYLF domain of BefA that reduced its membrane permeabilization activity. We also show that BefA can disseminate from the intestinal lumen via multiple anatomic routes to act directly on pancreatic tissue. Although small molecular metabolites of the gut microbiota are known to exert systemic effects on the host, especially in shaping immunity (Levy et al., 2016), the dissemination and functional impact of microbiota-derived proteins like BefA has not been described. Finally, we show that BefA’s β-cell expansion activity is conserved between zebrafish and mice and is dependent upon its membrane permeabilization activity, whereas the host-derived membrane permeabilizing Reg3 protein is sufficient to mimic BefA’s impact on islets. By investigating the structural and biochemical properties of BefA, our work reveals membrane permeabilization as a common mechanism of action by which both bacterial and host proteins stimulate pancreatic β-cell expansion.

## Results

### The structure of BefA reveals the SYLF domain that confers its function

To better understand how BefA functions, we determined its 3D atomic structure to 1.3 Å resolution, revealing a compact partial β-barrel carboxy-terminal domain with three flanking α-helices and five α-helices in the amino-terminal region (Figure 1A & B). The location of Helix 1 (H1) is likely constrained due to crystal packing contacts, and has been grayed out to show that the location in solution is unknown (Figure 1A-D). The high-resolution crystal structure of BefA confirms the backbone fold very recently reported for the solution NMR structure of a distantly related protein, BPSL1445 from *Burkholderia pseudomallei (Quilici et al., 2021)*, which taken together are the first representatives of a novel protein fold.

**Figure 1.**
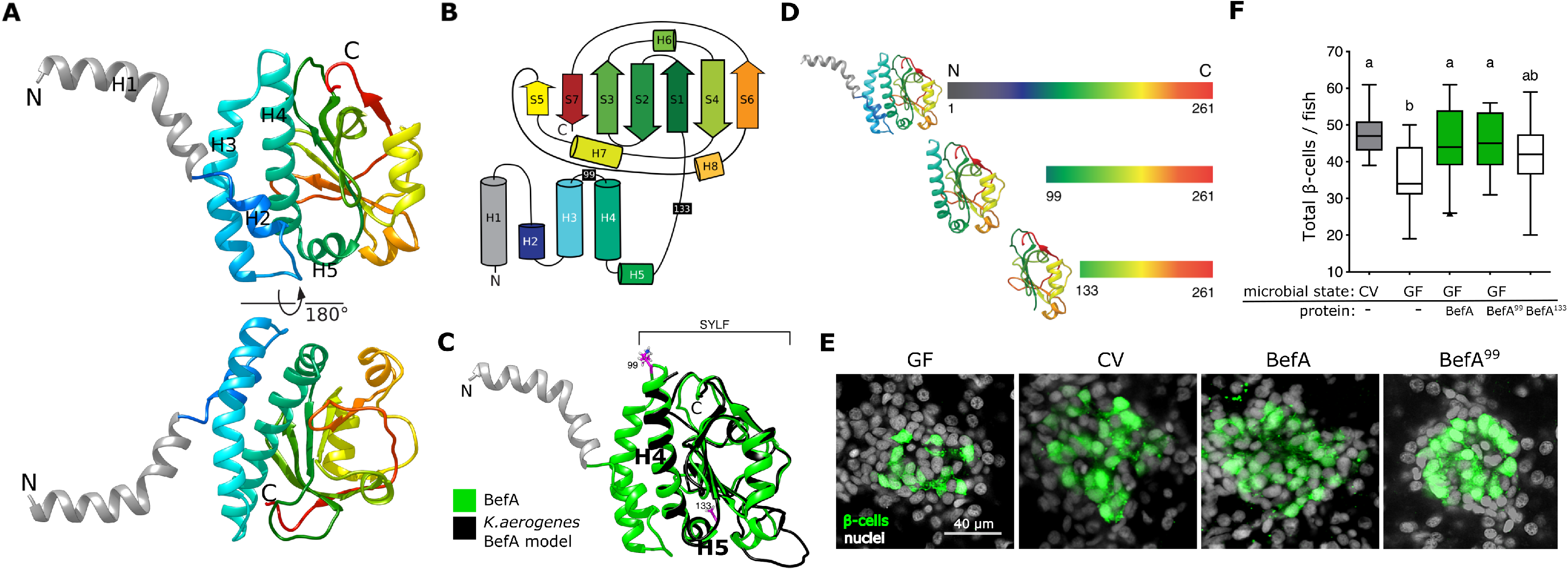
The structure of BefA reveals the SYLF domain that confers its function. (**A**) Ribbon diagram of the full length BefA protein in two orientations (rotated by 180° along the indicated axis) with the five amino terminal helices labelled H1-H5. H1 is colored gray to demonstrate that the position is constrained due to crystal packing. (**B**) 2D domain architecture map of BefA, to scale. Truncation locations from panel D are marked with corresponding amino-acid numbers (99 and 133) inside black rectangles. (**C**) The structure of BefA (green) overlaid with the predicted I-TASSER model of the structure of the *K. aerogenes* BefA homolog (black). Note that the amino terminus of the *K. aerogenes* model starts at H4 of the BefA structure. Residues 99 and 133 are shown as pink sticks in order to highlight that they denote the beginning of the BefA^99^ and BefA^133^ truncations (in panel D) respectively. (**D**) Scale schematics of the BefA truncation proteins, BefA^99^ and BefA^133^, accompanied by ribbon diagrams of their predicted structure. (**E**) Representative 2D slices from confocal scans through the primary islets of 6 dpf *ins:gfp* larvae which are either CV, GF, or GF treated with purified BefA or BefA^99^. Each slice is taken from the approximate center of the islet. Insulin promoter expressing β-cells are in green and nuclei are white. Scale bar = 40 μm. (**F**) Boxplots illustrating total β-cell quantifications from 6 dpf larvae in each treatment group, n≥15 larvae for each group from at least 2 experimental replicates. **In this, and in all subsequent figures containing boxplots**, CV data are colored grey, GF or control treatment groups are colored white, and BefA treated, or statistically similar groups, are colored green. **In all relevant panels and remaining figures**, box plot whiskers represent the 95% confidence interval of the data set **from pooled replicate experiments**. **In this and in all subsequent figures**, lowercase letters above groups indicate the results of post hoc means testing (Tukey) to denote statistically significant groupings.

To define the structural features corresponding to BefA’s β-cell expanding activity, we performed a structural comparison between the original *Aeromonas veronii* BefA and a distant BefA homolog identified in *Klebsiella aerogenes* (originally annotated as *Enterobacter aerogenes* (Hill et al., 2016)). Although the two amino acid sequences have only 34% identity (Figure S1A), the *K. aerogenes* homolog is sufficient to increase β-cell numbers in GF larval zebrafish (Hill et al., 2016). The structure of the *K. aerogenes* homolog was modeled using the Iterative Threading ASSEmbly Refinement (I-TASSER) server with BefA as a template (Krissinel and Henrick, 2004; Krissinel and Henrick, 2005; Roy et al., 2010; Yang et al., 2015b; Zhang, 2008). Overlaying the structure of BefA and the model of the BefA homolog revealed that despite the low amino acid sequence conservation between these two proteins, they are predicted to share a high degree of structural similarity starting at residue 99 at the beginning of H4 (RMSD = 0.84 Å) (Figure 1C). The absence of the three N-terminal α-helices (H1-H3) from the shorter *K. aerogenes* homolog model suggests that these helices are dispensable for function. Alignment with the NMR structure for BPSL1445 (PBD ID: 7OFN), a protein of unknown function from *B. pseudomallei* that shares 22% sequence identity with BefA, showed a common backbone fold across the SYLF domain with an RMSD value of 1.8 Å (Figure S1B). Additionally, we selected two well-studied eukaryotic SYLF-containing proteins from yeast (Ysc84) and humans (SH3YL1) and again used I-TASSER to predict the 3D structures of their SYLF domains. Both eukaryotic models are predicted to superimpose closely with our proposed BefA SYLF domain (RMSD values of 1.62 Å for Ysc84 and 1.92 Å for SH3YL1), beginning at residue 99 (Figure S1C), suggesting a high degree of structural conservation in SYLF domains across kingdoms. The Conserved Domain Databank (Marchler-Bauer et al., 2017) defines the SYLF domain in BefA as starting at residue 114, midway through H4 (Figure S1A, magenta arrowhead), but our structural analysis suggests that the conserved SYLF domain includes the entire helix, starting from residue 99 in BefA (Figure 1C).

With these new insights into the structure of the SYLF domain and the availability of many new bacterial genome and metagenome sequences, we expanded on our homology searches for bacterial BefA homologs (Hill et al., 2016), focusing on the SYLF sequence. Genes encoding SYLF-containing proteins were found in both environmental and host-associated bacteria across a variety of aquatic and terrestrial environments (Figure S1C). Our analysis identified many more examples of homologs like the *K. aerogenes* BefA and the *B. pseudomallei* BPSL1445, consisting almost entirely of the SYLF domain. Indeed, a survey of the average amino acid length of prokaryotic and eukaryotic SYLF-containing proteins shows the typical prokaryotic ones to be approximately the length of the 195 aa SYLF domain, whereas the eukaryotic proteins are significantly longer (Figure S1E), being comprised of one or more additional domains.

We predicted that the SYLF domain of BefA would be sufficient to mediate the pro-proliferative effect on larval zebrafish β-cells. To test this, we cloned two truncated BefA proteins. One truncation comprised amino acids 99 – 261, which encode the entire SYLF region by our definition (BefA^99^) (Figure 1D). The second shorter truncation incorporated amino acids 133-261, which correspond to the C-terminus compact partial β-barrel with three flanking α-helices (BefA^133^) (Figure 1D). We tested each purified truncated BefA protein for the ability to expand β-cell mass when incubated with GF *Tg(−1.0insulin:eGFP)* (*ins:gfp*) larvae from 4 to 6 days post fertilization (dpf). Quantifications of the total β-cell numbers in these fish confirm that BefA^99^ was equally effective at rescuing GF β-cell expansion as full length BefA (Figure 1E & F), whereas the BefA^133^ truncation only conferred a partial rescue of GF β-cell numbers (Figure 1F). Collectively, these results illustrate that the functional region of the BefA protein for inducing β-cell expansion is contained within the C-terminal SYLF domain.

### BefA induces membrane permeabilization

While SYLF domains are found across diverse forms of life, their mechanisms of action have only been investigated in a few proteins. Studies in yeast, plants, and mammals suggest that the SYLF domain commonly drives interactions at lipid membranes, helping to orchestrate complex biological processes such as motility and endocytosis (Urbanek et al., 2015b). Previously published evidence suggests that SYLF domain containing proteins mediate these important events by binding to membrane lipids (Hasegawa et al., 2011; Quilici et al., 2021; Sutipatanasomboon et al., 2017; Urbanek et al., 2015b) and/or actin filaments (Robertson et al., 2009b; Urbanek et al., 2015b). Therefore, we looked for biochemical interactions between BefA and lipids or F-actin.

The yeast protein Ysc84p contains actin binding function in its N-terminal SYLF domain (Hasegawa et al., 2011; Robertson et al., 2009b) and some of these actin binding residues are conserved in BefA. Therefore, we performed an F-actin co-sedimentation assay. Unlike our positive control, cortactin, we could not detect any significant enrichment of either BefA or BefA^99^ in the presence of F-actin (Figure S2A), indicating that BefA does not share Ysc84p’s ability to bind F-actin.

Since several other SYLF domain containing proteins can bind to phosphatidyl inositols (PIPs) (Hasegawa et al., 2011; Quilici et al., 2021; Sutipatanasomboon et al., 2017; Urbanek et al., 2015b), we next investigated the potential lipid and membrane binding activities of BefA. We used commercial lipid strip assays, which indicated no binding of BefA to PIPs, but consistent binding to phosphatidyl serine (PS) and cardiolipin, both negatively charged lipids (Figure S2B). The addition of calcium, which commonly mediates binding to negatively charged lipids, did not result in additional binding activity or increased affinity (Figure S2B). Several lipid binding residues previously identified in eukaryotic SYLF domains are conserved in BefA. To test whether any of these residues are necessary for BefA to bind to lipids, we created point mutations in BefA at several of these sites. We noted that several of these point mutants appeared to reduce BefA’s affinity for PS (Figure S2B), although conclusions from this assay are limited by the fact that these strips do not display lipids in the biologically relevant organization of membrane bilayers.

To test BefA’s interactions with lipid bilayers, we incubated either BefA or BefA^99^ with synthetic Texas Red-labelled giant lipid vesicles (Veatch, 2007) (Figure 2A). In the presence of BefA or BefA^99^ we observed a high frequency of large vesicles joined to smaller vesicles, a morphology we refer to as multi-vesicular vesicles (Figure 2A & B). We observed similar vesiculation with both neutral and negatively charged vesicles containing PS. We also noted that BefA treatment resulted in an increase in the total concentration of single and multi-vesiculated vesicles (Figure 2C & S2C), suggesting that multi-vesiculated vesicles are formed by the budding and fragmentation of larger vesicles, as schematized in Figure 2D.

**Figure 2.**
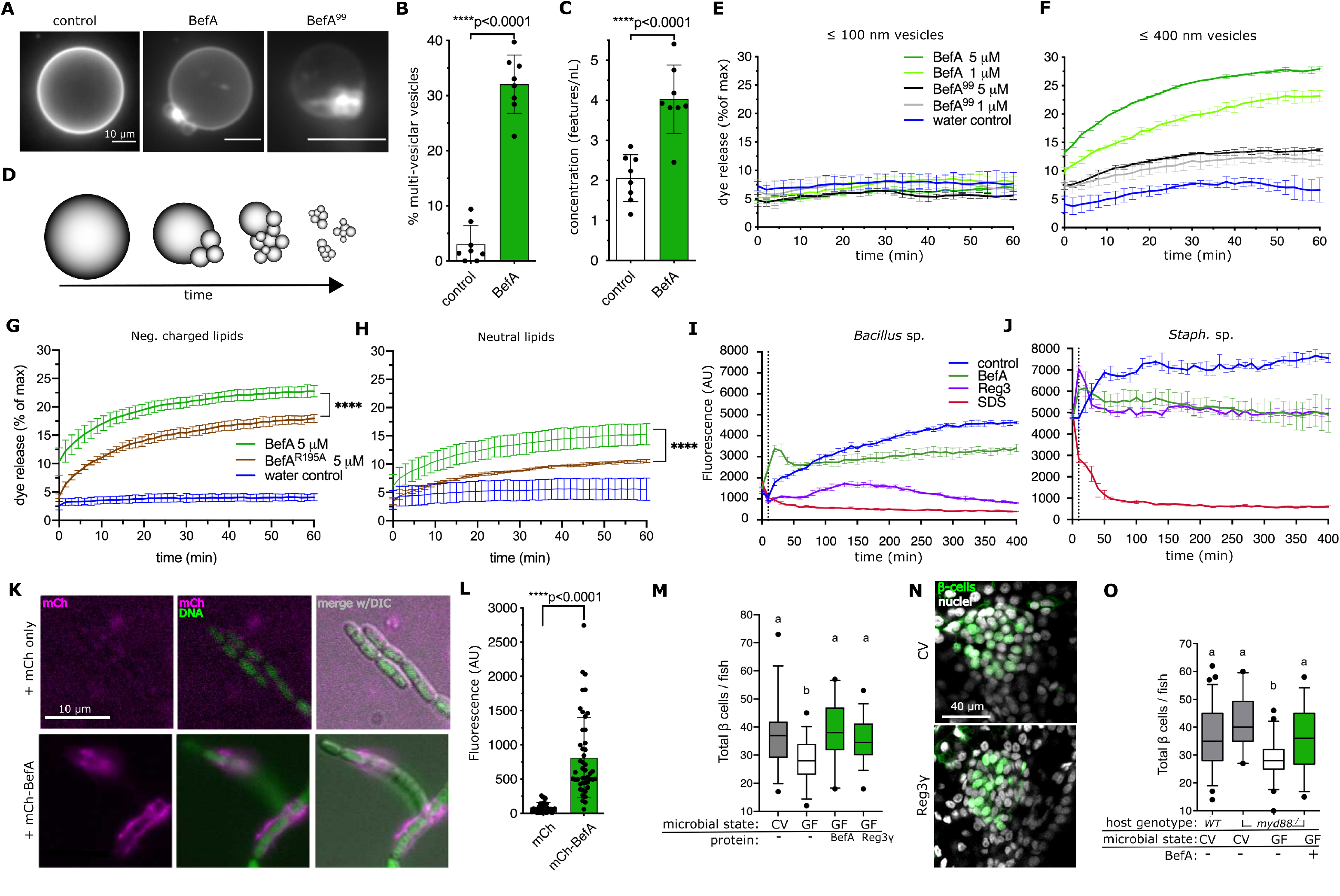
BefA induces membrane permeabilization. **(A)** Neutral (DOPC), fluorescently labeled vesicles in a range of sizes were treated with water (representative vesicle on left), 2 μM BefA (representative vesicle center), or 9 μM BefA^99^ (representative vesicle right) and imaged using light sheet fluorescence microscopy. All scale bars = 10 μm. **(B)** Model showing predicted vesiculation and subsequent fragmentation over time upon BefA treatment. **(C)** Each dot represents an image stack at either 35 or 50 min after treatment with 2 µM BefA, 32-150 vesicles per stack. (n = 8 stacks and total vesicles quantified ≥ 457 per treatment) Error bars represent the standard deviation from the mean. **(D)** The concentration of treated or untreated vesicles was quantified by summing the number of single plus multi-vesiculated vesicles (features) per volume for each image stack. (n = 8 stacks). Error bars represent the standard deviation from the mean. **(E&F)** Neutral, extruded vesicles were loaded with quenched dye and treated with water (control, blue), BefA at 1 and 5 μM (green traces), or BefA^99^ at 1 and 5 μM (gray traces). Leakiness of vesicles over time was monitored by release of dye as a percentage of maximal dye release upon addition of detergent. Vesicle sizes for (**E**) were less than or equal to 100 nm in diameter or less than or equal to 400 nm for (**F**). Lines follow the mean of duplicate experiments with error bars denoting the standard deviation of the mean between experiments for each time point. (**F**) Two-way ANOVA between treatment groups p<0.0001. **(G&H)** Negatively charged vesicles containing PS (**G**) or neutral vesicles (**H**) were loaded with quenched dye and treated with water (control, blue), BefA at 5 μM (green traces), or BefA^R195A^ at 5 μM (brown traces). Leakiness of vesicles over time was monitored by release of dye as a percentage of maximal dye release upon addition of detergent. Vesicles were less than or equal to 400 nm in size. Lines follow the mean of duplicate or triplicate experiments with error bars denoting the standard deviation at each time point. ****p value<0.0001 based on Two-Way ANOVA multiple comparisons test. **(I&J)** Bacterial cell permeabilization was monitored for *Bacillus* sp. and *Staphylococcus* sp. using CyQUANT Direct Red plate assay for negative control (black), 7 μM BefA (green) and 3 μM Reg3α (purple) treated cells. 0.08% SDS detergent (red) was used as a positive control. In contrast to panels E&F, permeabilization is detected as dye released from cells becomes quenched, resulting in decreasing fluorescent signal. Lines follow the mean of 3 replicate experiments with error bars denoting the min and max spread of the data between experiments for each time point. **(K)** Representative images of *Bacillus* sp. incubated with mCh only (top) or mCh-BefA (bottom). Magenta = mCh protein, green = DNA stained with SYBRGreen, DIC = differential interference contrast microscopy. Scale bar = 10 μm. **(L)** Quantification of BefA binding to *Bacillus* sp. after incubation with mCh-BefA or control mCh alone. The mean fluorescence intensity of the mCh channel along the length of each cell was measured and graphed. (n≥33 individual bacterial cells per treatment) Error bars represent the standard deviation from the mean. (**M**) Boxplots illustrating total β-cell quantifications from 6 dpf larvae in each treatment group, n≥25 larvae for each group. (**N**) Representative 2D slices from confocal scans through the primary islets of 6 dpf *ins:gfp* larvae which are either CV, or Reg3g treated. Each slice is taken from the approximate center of the islet. Insulin promoter expressing β-cells are in green and nuclei are white. Scale bar = 40 μm. (**O**) Boxplots illustrating total β-cell quantifications from 6 dpf larvae in each treatment group, n≥19 larvae for each group.

Induction of membrane curvature, which occurs in the formation of multi-vesicular vesicles, is a common characteristic of membrane permeabilizing antimicrobial proteins (Guha et al., 2019). To determine whether BefA could impair membrane integrity, we performed a dye release assay on vesicles encapsulating fluorescent dye. While little to no dye was released by smaller vesicles (≤100 nm in diameter) upon treatment with either BefA, BefA^99^, or a water control (Figure 2E), a significant amount of dye was released in a concentration dependent manner when BefA or BefA^99^ was added to larger vesicles (≤400 nm in diameter) (Figure 2F), suggesting BefA may have greater affinity for lower membrane curvature. We next tested whether a BefA mutant with reduced PS binding in the lipid strip assay would have reduced vesicle permeabilizing activity. We found that BefA^R195A^ released significantly less dye from both neutral and negatively charged PS-containing vesicles compared to wild type BefA (Figure 2G & H). Arginine 195 is located on the same protein face immediately adjacent to a stretch of residues shown in BPSL1445 to interact in solution with phospholipids (Quilici et al., 2021).

Many antimicrobial proteins target bacterial cell membranes by inducing membrane curvature (Mukherjee and Hooper, 2015). To assess whether BefA could compromise bacterial membrane integrity, we used the CyQUANT Direct Red cell permeability assay, whereby cell leakiness is measured as quenching of a membrane permeable fluorescent DNA dye by a cell-impermeant suppressor. We tested two novel Gram-positive isolates from zebrafish belonging to the genera *Bacillus* and *Staphylococcus.* These cells were readily permeabilized by the detergent SDS, as well as by the well-characterized pore-forming antimicrobial protein, Reg3α (Figure 2I & J), which is secreted by host enterocytes to maintain barrier function against the gut microbiota (Vaishnava et al., 2011). BefA also induced bacterial membrane permeabilization in *Bacillus* and *Staphylococcus* with levels of permeabilization comparable to that of Reg3α in the case of *Staphylococcus* (Figure 2I & J). We also tested whether BefA bound directly to bacterial cells, using an mCherry (mCh)-BefA fusion protein. We observed clear binding of mCh-BefA to the outside of the *Bacillus sp.* and very little to no binding of the unconjugated mCh protein (Figure 2K & L). BefA binding appeared to be enriched at junctions between cells, where nascent cell wall synthesis during septation may allow greater access to the cell membrane (Pasquina-Lemonche et al., 2020).

We speculated that BefA’s capacity to permeabilize membranes was the basis for its β-cell expanding activity. If this were the case, then other unrelated membrane-permeabilizing proteins should also be able to increase β-cell mass. To test this, we exposed GF larvae to purified Reg3γ, the murine homolog of human Reg3α. The addition of Reg3γ was sufficient to rescue β-cell numbers to that of CV or BefA treated levels (Figure 2M). Overall morphology of larval islets treated with Reg3γ was similar to that of CV, suggesting that addition of the AMP did not cause significant injury or damage to the developing cells (Figure 2N).

Host AMPs are often produced downstream of Toll-like receptor (TLR) activation signaling cascades. Both pancreatic β-cells and intestinal cells express TLRs, which recognize microbial products (Garay-Malpartida et al., 2011; Vives-Pi et al., 2003), making TLR signaling a candidate mechanism to explain both BefA and Reg3γ activity on β-cells. To test whether TLR signaling was required for BefA sensing, we quantified β-cell numbers in zebrafish larvae carrying a mutation in the *myd88* gene (Burns et al., 2017). Myd88 is a universal adaptor protein downstream of most TLR and IL1R activation. We did not observe any difference in the number of β-cells between wild-type (WT) and *myd88^-/-^* larvae that were raised conventionally (Figure 2O). Furthermore, treatment of GF *myd88^-/-^* larvae with BefA resulted in an increase of β-cells to numbers similar to CV fish (Figure 2O). Together these results suggest that TLR signaling is not required for BefA sensing.

### BefA interacts directly with pancreatic β-cells

Our finding that BefA and another membrane-permeabilizing protein could expand pancreatic β-cells suggested that BefA’s effects would be mediated through direct interaction with target host cells. To visualize interactions between BefA’s SYLF domain and host cells, we utilized fluorescently tagged versions of BefA^99^ (mCh- or mNeonGreen (mNG)-BefA^99^), which we showed induced a similar increase in β-cell numbers in GF larvae as the untagged full length BefA; unconjugated fluorescent proteins did not increase β-cell number (Figure 3A). First, we added mNG-BefA^99^ to dissociated zebrafish cells cultured from dissected intestines of larval zebrafish. These preparations contained insulin positive pancreatic β-cells and a variable mixture of cells from other digestive tract tissues. While we detected little to no mNG signal in cultures incubated with mNG alone, we saw distinct punctate labeling of mNG-BefA^99^ co-localized with β-cells (Figure 3B, yellow arrowheads) as well as other non-insulin expressing cells (Figure 3B, white arrowheads). We found that an average of 21 (std. dev. = 14) percent of β-cells associated with mNG-BefA^99^, whereas only 1 (std. dev. = 3) percent of β-cells associated with mNG alone, demonstrating that the SYLF domain of BefA is sufficient to facilitate a direct interaction with β-cells. To determine the temporal dynamics of this interaction, we performed a longitudinal imaging experiment. Dissociated larval islet cells were treated with mNG-BefA^99^ and imaged using confocal microscopy for a period of 30 minutes. A subset of cells in the islet displayed an initial uniform surface fluorescence with mNG-BefA99, followed by the appearance of surface puncta as early as 10 minutes post-exposure (Figure S3A). After 20 minutes, the puncta appear to become internalized into the cells (Figure S3A), consistent with a process of protein endocytosis.

**Figure 3.**
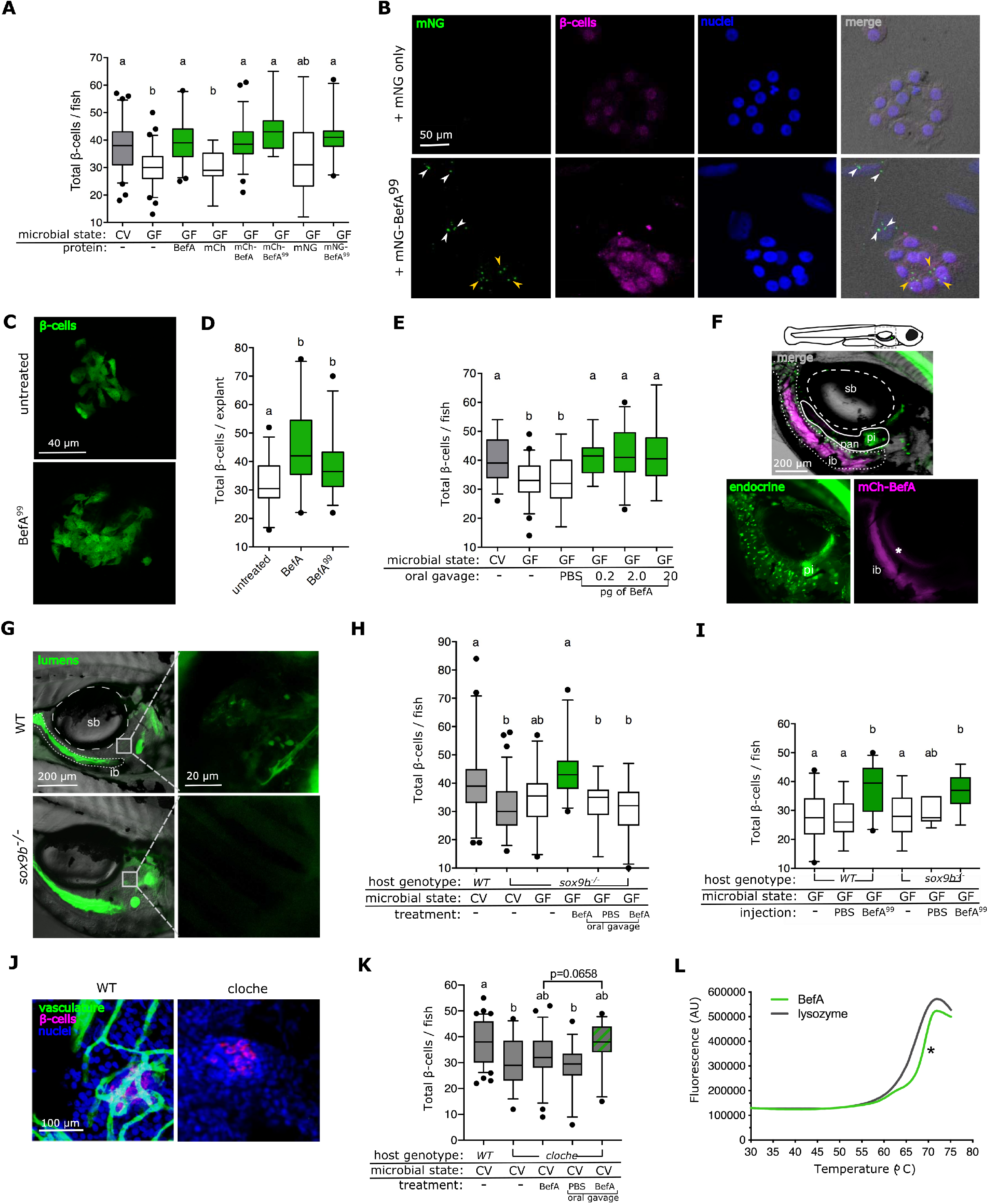
BefA interacts directly with host β-cells. (**A**) Boxplots illustrating total β-cell quantifications from 6 dpf larvae in each treatment group, n≥10 larvae for each group. (**B**) Confocal images of dissociated larval gastrointestinal cell cultures, treated with either mNG or mNG-BefA^99^. Panels from left to right: Insulin in magenta, mNG in green, nuclei in blue, and DIC overlay with fluorescent channels merged. White arrowheads indicate some of the mNG puncta associated with non-insulin expressing cells. Yellow arrowheads indicate some of the mNG puncta associated with β-cells. Scale bar = 50 μm. (**C**) Representative 2D slices from confocal scans through islet explants after treatment. Insulin promoter expressing β-cells are in green. Scale bar = 40 μm. (**D**) Boxplots illustrating total β-cell quantifications from primary islet explants in each treatment group, n≥20 explants for each group. (**E**) Boxplots illustrating total β-cell quantifications from 6 dpf larvae treated by oral gavage with varying concentrations of BefA (in picograms (pg)), n≥18 larvae for each group. (**F&G**) Cartoon larvae aged 4 dpf with gray dotted outline denoting the trunk region imaged. For orientation, the swim bladder (sb) is outlined in large white dashed lines, the intestinal lumen is outlined in small white dashed lines, and the pancreas (pan) is outlined in a solid white line, ib = intestinal bulb, pi = primary islet. (**F**) Top panel = merge of bottom two panels with DIC overlay. Lower left panel = *Nkx2.2* promoter expressing endocrine tissue in green. Lower right panel = mCh-BefA in magenta. * = reflection of mCh signal off swim bladder. (**G**) Left panels represent wild-type (top) and *sox9b^-/-^* (bottom) larvae fed BODIPY FL-C_2_ (green), insulin-expressing cells = magenta, DIC overlay = grey. Scale bar = 200 μm. White box around the primary islet indicates the region of the zoom inset in right panels. Scale bar = 20 μm. (**H&I**) Boxplots illustrating total β-cell quantifications from 6 dpf larvae in each treatment group, (**H**) n≥17 or (**I**) n≥6 larvae for each group. (**J**) Representative confocal scans through primary islets of WT and *cloche* mutant larvae. Insulin promoter expressing β-cells are in magenta, vasculature (anti-kdrl) staining in green, nuclei in blue, scale bar = 100 μm. (**K**) Boxplots illustrating total β-cell quantifications from 6 dpf larvae in each treatment group, n≥15. (**L**) The thermal stability of BefA (green trace) and lysozyme (gray trace), measure by the Thermofluor assay. *T_m_ value of BefA as indicated by the inflection point of the curve = 68° C.

We next tested if direct interaction with BefA is sufficient to exert a proliferative effect on larval β-cells using primary zebrafish islet explant cultures. We micro-dissected the primary islet from GF 4 dpf *ins:gfp* larvae (Figure S3B) and maintained them *ex vivo* in sterile cell culture media for 48 hours untreated or in the presence of full length BefA or BefA^99^. *Ex vivo* larval β-cells appeared healthy after 48 hours as indicated by robust insulin promoter driven *gfp* transgene expression, absence of TO-PRO-3 Iodide (TOPRO) incorporation marking dead or dying cells, and an otherwise normal appearance (Figure S3C). After the treatment period, we observed significantly more β-cells in explants treated with either BefA or BefA^99^ than in those that received only rich cell culture medium (Figure 3C & D), showing that BefA can elicit β-cell expansion via a direct interaction, and that the SYLF domain is sufficient to mediate this effect.

After finding BefA’s direct effect on pancreatic cells, we sought to determine whether this protein, secreted by gut bacteria, could disseminate to the pancreas. Plausible routes for dissemination from the gut lumen would be via the vasculature or through reflux up the hepatopancreatic duct (HPD), which connects the liver, the gallbladder, and the pancreas to the gut for the passage of bile and digestive enzymes (Field et al., 2003). Additionally, because BefA could elicit β-cell expansion when added to the water column, we considered the possibility that it could be absorbed through the skin and gills into the bloodstream. To confirm that gut lumenal BefA could affect pancreatic β-cells, we used oral microgavage to deliver different doses of BefA directly into the intestinal lumen of 4 dpf larval zebrafish, before quantifying cell numbers at 6 dpf. We found that administration of BefA directly into the gut lumen was sufficient to increase β-cell numbers in GF fish compared to the gavage vehicle control of PBS, even at the lowest dose tested of 0.2 pg (Figure 3E).

We next sought to visualize systemic BefA dissemination following oral microgavage of *Tg*(*Nkx2.2*:*gpf*) GF larvae, which express GFP in intestinal and pancreatic endocrine tissues, allowing us to visualize both intestinal and pancreatic anatomical landmarks. Using either mCh-BefA or mNG-BefA and corresponding mCh or mNG control proteins, fluorescence was readily visualized within the intestinal lumen (Figure 3F) and in lysosome rich enterocytes (LREs) (Figure S3D) that have recently been described as a site of active, non-selective protein internalization (Park et al., 2019). Given the strong lumenal and LRE fluorescent signals and our finding that even very low doses of BefA will induce β-cell expansion, we concluded that visualizing functionally relevant, BefA-specific host cell interactions was beyond our *in vivo* imaging capabilities. We next explored BefA’s routes of dissemination using zebrafish mutants.

To test the possibility of BefA reflux through the ductal network connecting the intestine and pancreas, we used *sox9b^-/-^* mutant zebrafish in which HPD development is compromised (Delous et al., 2012; Manfroid et al., 2012) such that bile secretion into the intestine is blocked in adulthood (Delous et al., 2012). We tested the patency of the HPD in *sox9b^-/-^*4 dpf larval zebrafish using the short chain fluorescent lipid analog, BODIPY-FL C_2,_ which fills lumenal spaces of the GI tract because it is not appreciably metabolized (Carten et al., 2011). After feeding BODIPY-FL C_2_ to WT larvae, it could be detected in intrapancreatic lumenal structures likely to be the intra-pancreatic ducts (IPD), which are contiguous with the intestinal lumen (Figure 3G), whereas in *sox9b^-/-^* larvae, little to no fluorescent signal from BODIPY-FL C_2_ could be found in the pancreas despite its obvious ingestion and presence in the intestinal lumen (Figure 3G). We next tested whether this loss of patency through the HPD in *sox9b^-/-^* mutant larvae would affect the ability of either the microbiota or BefA to influence the growth of the larval β-cell population. CV *sox9b^-/-^* larval β-cell numbers were significantly reduced compared to CV *WT* animals, and were equivalent to GF *sox9b^-/-^* larvae (Figure 3H), consistent with *sox9b^-/-^*larvae being unable to respond to BefA from their endogenous gut microbiota. However, GF *sox9b^-/-^* larvae that were treated with BefA in their water had significantly increased β-cell numbers (Figure 3H), indicating that *sox9b^-/-^*larvae were competent to respond to BefA and suggesting that BefA added to the water column may act through a different route than BefA produced by gut bacteria. To test this hypothesis, we delivered a high dose of BefA via oral microgavage to 4 dpf GF *sox9b^-/-^* larvae and quantified β-cell numbers at 6 dpf. When delivered directly to the intestinal lumen, BefA was not capable of increasing β-cell numbers in *sox9b^-/-^* larvae (Figure 3H). Together, these results suggest that patency of the HPD is required for delivery of BefA from the gut lumen, but that BefA is also able to reach its target cell population through an alternative route.

To test whether the vasculature was an alternative route through which BefA could disseminate, we injected BefA^99^ into the region of the developing circulation valley of GF larval zebrafish. Larvae injected with BefA^99^ had significantly greater β-cell numbers compared to an injection control (Figure 3I). We next injected BefA into *sox9b^-/-^* larvae and found a significant increase in β-cells (Figure 3I). Together, these results suggest that BefA added to the water column is absorbed and disseminated through the blood stream to elicit β-cell growth in *sox9b^-/-^* larvae.

To further investigate the role of the vasculature in BefA dissemination, we utilized *cloche* mutant zebrafish which have grossly abnormal hematopoietic development and lack blood vessels, resulting in severe edema and early lethality (Reischauer et al., 2016; Stainier et al., 1995) (Figure 3J). Because of these defects and low yields of offspring, *cloche* mutants were not amenable to our GF derivation protocols and we were limited to experiments in CV larvae. CV *cloche* mutant larvae had significantly reduced numbers of β-cells compared to their CV *WT* clutch-mates (Figure 3K), and addition of a concentrated dose of BefA to the water column was not sufficient to fully rescue this effect. However, we were able to increase β-cell numbers more if we delivered a high dose of BefA to the *cloche* mutants via oral microgavage (Figure 3K), suggesting that these mutants are still competent to respond to BefA via the HPD, but fail to do so when BefA is produced in lesser amounts by endogenous gut microbiota or added in the water column. Collectively, these data support the idea that secreted BefA can reach distant β-cells in the pancreas both by traveling from the gut lumen through the HPD or by being absorbed into the vasculature. Furthermore, these results suggest that access to the pancreas is required for BefA to exert its function on β-cells, supporting the hypothesis that it acts in a direct manner *in vivo*.

The intestinal lumen, the HPD, and the blood stream represent drastically different chemical microenvironments that could influence protein folding. When purifying BefA protein, we noted that it was soluble in pure water without ions or buffer at concentrations exceeding 20 mg/mL, indicating high structural stability. Using a thermal stability assay, we determined BefA’s melting temperature to be approximately 68° C, comparable to the highly stable protein lysozyme, which melts at 65° C (Ku et al., 2009) (Figure 3L), suggesting that BefA’s structure would be maintained while disseminating through potentially harsh host environments.

### BefA’s direct activity on host β-cells is conserved in mice

Our analysis of SYLF domain encoding genes uncovered multiple homologs in *Enterobacteriaceae* members of mammalian microbiotas (Figure S1C), suggesting the possibility that the mammalian pancreas is exposed to BefA homologs during neonatal β-cell development. Therefore, we sought to determine whether the effects of the microbiota and BefA on β-cell development were conserved in mammals. Like zebrafish, mice are born with a population of fully differentiated, insulin-secreting β-cells (Larsen and Grapin-Botton, 2017). During neonatal life, which is comparable to the post-hatching age of zebrafish, mammalian β-cells undergo high levels of proliferation to establish a healthy amount of insulin-producing tissue capable of maintaining glucose homeostasis throughout life (Miller et al., 2009). To test for the ability of BefA to increase murine β-cell mass, we engineered the probiotic *E. coli* Nissle 1917 (Nissle) strain to express *befA* from a chromosomally integrated transgene. Importantly, wild-type Nissle (Nissle*^WT^*) has no endogenous copy of *befA* in its genome, it robustly colonizes both zebrafish and mice, and it is not pathogenic. Furthermore, since we found homologs of BefA within the genomes of other *E. coli* strains (Figure S1C) (Hill et al., 2016) we predicted that Nissle would be able to functionally produce and secrete BefA. To confirm this, we performed Western blots on bacterial cell free supernatants (CFS) using a polyclonal anti-BefA antibody (Figure S4A). CFS concentrated from wild-type *Aeromonas veronii* HM21 (*A. veronii*), the strain from which BefA was originally isolated, showed a strong signal at the expected size of BefA (29 kDa), while no signal was present in the lane containing CFS from our *befA* knockout *A. veronii* strain (Figure S4A), validating the specificity of the anti-BefA antibody. Similarly, we saw a strong band in the lane containing CFS from Nissle*^befA^* and no signal in the lane containing CFS from Nissle*^WT^*, indicating that our new transgenic Nissle strain was capable of secreting BefA protein (Figure S4A). To test the functionality of BefA protein secreted by Nissle*^befA^*, we mono-associated GF zebrafish larvae with either Nissle*^WT^* or Nissle*^befA^*. Larvae colonized with Nissle*^befA^* had significantly more β-cells than either GF larvae or those colonized with Nissle*^WT^* (Figure S4B), confirming that the BefA secreted by our transgenic strain was functional, and that the wild type strain was not capable of inducing β-cell expansion via an alternative mechanism.

We next examined postnatal β-cell development in wild type C57/BL6 mice with different microbial associations at postnatal day 12 (P12), just after neonatal β-cell proliferation levels have peaked (Miller et al., 2009). Importantly, mice of this young age are highly variable in overall weight due to differences such as litter size, which can drive similar differences in pancreas mass across our experimental groups (Figure S4C). Therefore, we quantified the average ratio of insulin-expressing area to total pancreatic cross-sectional area for each pup, rather than β-cell mass, which is the conventional unit in adult mice. We found that mice with a conventional or specific pathogen free (SPF) microbiota had significantly increased ratios of insulin-positive area compared to mice that were either GF or had been given antibiotics (ABX) from birth (Figure 4A & B), indicating that resident bacteria play a stimulatory role in mammalian β-cell development. To test whether BefA is sufficient to rescue reduced β-cell levels in mice lacking resident bacteria, we mono-associated GF mice with our Nissle strains by inoculating breeder pairs, and allowing them to transmit Nissle vertically to their offspring. As before, we analyzed β-cell area in P12 aged mice from each mono-association and found that pups colonized with Nissle*^befA^* had significantly higher insulin positive area than Nissle*^WT^* (Figure 4B & C). We also treated SPF pups concurrently with both antibiotics and purified BefA protein administered by oral gavage during postnatal life, and again analyzed insulin content of the pancreas. Purified BefA treatments were sufficient to rescue the total β-cell content of antibiotic-treated pups to levels similar to those of SPF (Figure 4A & B). We also allowed the mice from some of our Nissle mono-associated litters to grow to adulthood before we analyzed total β-cell mass. Like the pups, adult mice mono-associated with Nissle*^befA^* had significantly greater β-cell mass compared to those mono-associated with Nissle*^WT^* (Figure 4D). The overall pancreas mass of adult mice with these mono-associations was not different (Figure 4E), suggesting that mouse pancreas size is not affected by BefA. We also examined alpha-cell mass in adult GF and SPF mice, and found no significant differences (Figure S4D-F), similar to our previous findings in zebrafish (Hill et al., 2016). Collectively, these results demonstrate that BefA is capable of increasing the β-cell mass of mice, and that these effects are specific and long lasting.

**Figure 4.**
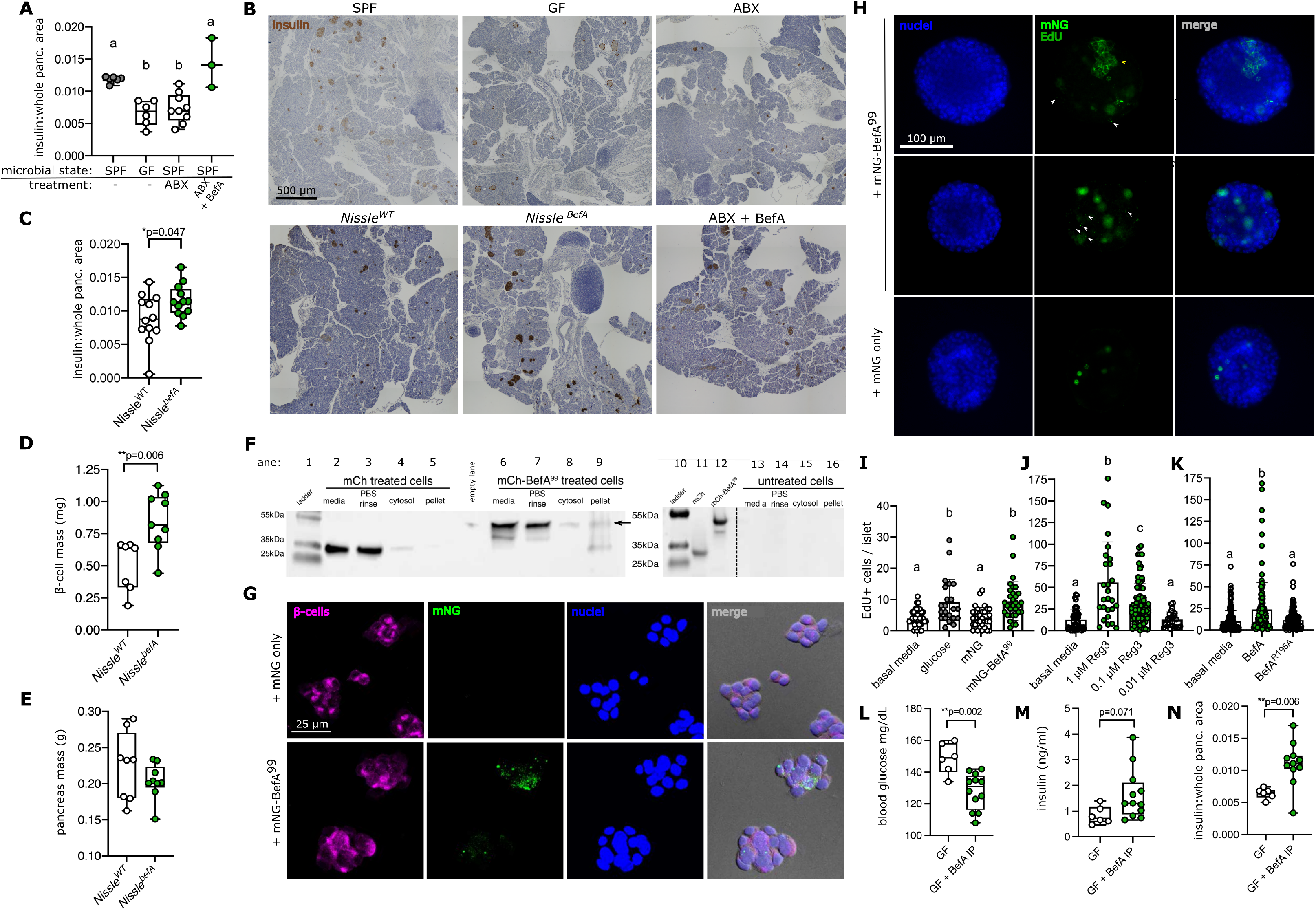
BefA’s direct activity on host beta-cells is conserved in mice. **(A**) Boxplots of quantifications of average ratio of insulin area to whole pancreas area across at least three cross sections per mouse pup aged P12. n≥5 mice in each treatment group except ABX + BefA where n=3. (**B**) Representative cross sections through postnatal day 12 (P12) mouse pancreata, ABX = Antibiotic treated, Hematoxylin staining nuclei (blue) and Insulin (brown). Because insulin staining in GF mice is faint, dashed brown outlines are meant to help distinguish insulin^+^ areas. Scale bar = 500 μm. **(C**) Boxplots of quantifications of average ratio of insulin area to whole pancreas area across at least three cross sections per mouse pup mono-associated with Nissle strains, aged P12. n=12 mice in each treatment group. *p value for Student’s T-test. (**D**) Boxplot illustrating total β-cell mass in milligrams (mg) of adult mice, mono-associated with Nissle strains, n≥8 for each group, *p value for Student’s T-test. (**E**) Boxplot illustrating total pancreas mass in grams (g) in adult mice mono-associated with Nissle strains, n≥8 for each group. (**F**) Western blots for mCherry on βTC-6 cell cultures treated with either mCh (29 kDa) or mCh-BefA^99^ (45 kDa) proteins. Lane 1 & 10: ladder. Lanes 2-5: mCherry treated culture components. Lanes 6-9: BefA^99FL^ treated culture components. Lane 11: purified mCh protein. Lane 12: purified mCh-BefA protein. Lanes 13-16: untreated culture components. Arrow: mCh-BefA^99^ band within cell pellet fraction. (**G**) Confocal images of βTC-6 treated with either mNG or mNG-BefA^99^. Panels from left to right: Insulin in magenta, mNG in green, DAPI in blue, and DIC overlay with fluorescent channels merged. Scale bar = 25 μm. (**H**) Fluorescent images of primary mouse islets from P12 SPF pups, after 48-hour treatment with either mNG-BefA^99^ (top and middle rows), or mNG only/unconjugated (bottom row). Left column = DAPI stain for nuclei in blue, middle column = mNG (either punctate as pointed out by white arrowheads and outer membrane associated as denoted by yellow arrowhead) and EdU (nuclear) in green, right column = merge of blue and green channels. Scale bar = 100 μm. (**I-K**) Bar graphs illustrating mean EdU labelled cells in each primary islet per treatment group, each dot represents one islet, whiskers denote std. dev. n≥22 islets. (**L**) Blood glucose readings (mg/dL) from nonfasted mice of each treatment group. (**M**) Serum insulin measurements from nonfasted mice of each treatment group. (**N**) Boxplots of quantifications of average ratios of insulin area to whole pancreas area across at least two cross sections per mouse. (**L-N**) n≥6 mice in each treatment group, aged P21. *p value for Student’s T-test.

We next performed several experiments to determine if BefA acts directly on murine β-cells. First, we exposed cultured βTC-6 cells, an immortalized murine β-cell line (Poitout et al., 1995), to mCh alone or mCh-BefA^99^. Following an incubation of several hours, βTC-6 cells were washed, and fractions of the cell membranes, cytosol, and rinse were analyzed by Western blot. A band at the expected size of mCh-BefA^99^ (45 kDa) localized to all the βTC-6 culture components, including the cell membrane fraction (Figure 4F), whereas bands of the expected size for mCh alone (29 kDa) were not found in the cell membrane pellet (Figure 4F). We next imaged mouse βTC-6 cells incubated with our brighter mNG-BefA^99^ protein. Similarly to our observations with dissociated zebrafish cells (Figure 3B, S3A), we detected little to no mNG puncta associated with β-cells in cultures incubated with mNG alone (mean percentage of cells co-localized with mNG = 0.2, std. dev = 0.6), but observed significant punctate labeling of mNG-BefA^99^ co-localized with β-cells (mean percentage of cells co-localized with mNG = 11, std. dev = 5) (Figure 4G), indicating that BefA is capable of directly interacting with cells of murine origin, and that the SYLF domain is sufficient for this activity.

We next tested whether direct interaction of BefA’s SYLF domain with murine pancreas tissue would increase β-cell proliferation. Primary mouse islets were isolated from wild type SPF Swiss Webster mice aged P12 and treated *in vitro* with either mNG-BefA^99^, mNG alone, 10 μM glucose, or were left untreated for 48 hours. To measure islet cell proliferation, we treated with the nucleotide analog EdU during the final 4-6 hours of incubation. Islets treated with mNG-BefA^99^ showed distinct BefA^99^ puncta, similar to the cultured βTC-6 cells and absent from mNG treated islets (Figure 4H). The addition of either mNG-BefA^99^ or our positive control, glucose, resulted in significantly higher levels of EdU incorporation per islet than either untreated or mNG treated groups (Figure 4I), indicating that the SYLF domain of BefA is sufficient to elicit proliferation of neonatal β-cells in mice.

To determine whether membrane permeabilization is sufficient to elicit mammalian islet cell turnover, we next treated primary neonatal mouse islets with the pore-forming protein, Reg3α. Reg3α treatment resulted in a significant dose-dependent increase in islet cell proliferation as measured by EdU incorporation (Figure 4J). To test whether membrane permeabilization was necessary for BefA-mediated islet cell turnover, we treated primary neonatal mouse islets with the BefA^R195A^ mutant that permeabilizes lipid vesicles less effectively (Figure 2G & H). When compared to full-length wild type BefA, BefA^R195A^ elicited significantly reduced levels of EdU incorporation (Figure 4K). Taken together, our results indicate that BefA elicits β-cell proliferation via a mechanism of increasing membrane permeabilization.

Lastly, to test whether BefA could disseminate through alternate anatomical routes to elicit a response from β-cells *in vivo*, as we had observed in our zebrafish model, we injected purified BefA protein intraperitoneally (IP) into GF Swiss Webster neonatal mice and analyzed β-cell mass at weaning (P21). Blood glucose measurements taken from mice injected with BefA were significantly lower than untreated mice (Figure 4L), which was consistent with increased serum insulin levels (Figure 4M). Image analyses of the pancreata from these mice also revealed significantly higher levels of insulin expressing tissue in BefA injected samples (Figure 4N), indicating that IP delivery of BefA is sufficient to exert the same effect on the pancreas as when secreted by gut bacteria. Collectively, these data show that BefA is able to induce β-cell expansion in mice via multiple routes of delivery, highlighting the conserved capacity of a gut microbiota derived protein to disseminate through the body and impact the development of distant tissues.

## Discussion

The many bacterial genomes that make up the human microbiome encode a largely unexplored repertoire of diverse bioactive molecules that can impact animal biology. One such example is the secreted bacterial protein, BefA, which we showed can induce expansion of pancreatic β-cells (Hill et al., 2016). The novelty of such microbial products presents a unique challenge for uncovering their bioactive mechanisms. In the current study, we utilized a broad array of molecular, genetic and biochemical approaches to elucidate the mechanism of action of the BefA protein. Our findings reveal that BefA is a compact and stable membrane permeabilizing protein that directly stimulates β-cell expansion in vertebrates. We show BefA’s membrane permeabilizing activity is necessary to stimulate β-cell proliferation and that a non-homologous membrane permeabilizing protein, Reg3, recapitulates BefA’s impact on β-cells.

The BefA crystal structure reveals atomic features of the lipid-binding SYLF domain, which is widely distributed in predicted proteomes across the kingdom of life. While eukaryotic SYLF domain-containing proteins are typically multidomain proteins, the SYLF region comprises the bulk of BefA and its prokaryotic homologs. This suggests that the SYLF’s intrinsic membrane permeabilizing activity confers a selective advantage for bacteria, possibly as an antimicrobial weapon against competitors in multispecies communities (Granato et al., 2019) or as an agent of programmed cell death (Bayles, 2014). *befA* deficient *Aeromonas* mutants are at a slight competitive disadvantage in di-association with wild type *Aeromonas* (Hill et al., 2016) under conditions of normal host β-cell census, indicating that BefA benefits *Aeromonas* independently of bacterial competitors or insulin paucity. Although we do not yet fully understand BefA’s role in *Aeromonas* fitness, our studies reveal how the protein’s biochemical properties of stability and membrane permeabilizing activity mediate its impact on host biology.

BefA’s compact and stable structure likely endows it with the capacity to persist in diverse extra-cellular environments. Our finding that BefA can directly induce β-cell proliferation in primary islet cultures suggests that it disseminates to the pancreas from the intestinal lumen and water column. This conclusion is supported by imaging of fluorescently labelled BefA and functional characterization of BefA through multiple routes of administration. Furthermore, the BefA recalcitrance of larval mutants lacking a patent HPD or vasculature indicates that both of these routes are involved in the pancreatic response to BefA. Reflux up the HPD has long been suspected as a potential route of bacterial transmission to both the pancreas and the liver, where bacteria have been found to drive carcinogenesis (del Castillo et al., 2019; Pushalkar et al., 2018; Riquelme et al., 2019; Tilg et al., 2016). In addition, Zhang and colleagues recently highlighted the bloodstream as a mode of delivery for bacterial products directly to the pancreas. Their study showed that microbial ligands of the Nod1 receptor could access the blood stream in order to subsequently bind to β-cells and promote insulin trafficking (Zhang et al., 2019). Together with our findings, these results further our mechanistic understanding of how microbial products from the gut impact distant host tissue biology.

Our studies in simplified synthetic lipid vesicle systems revealed that BefA and its SYLF domain have intrinsic membrane vesiculating and permeabilizing activity. Membrane permeabilizing proteins and peptides are produced across the kingdom of life, from viral encoded viroporins to animal antimicrobial proteins. Many of these proteins act by changing membrane curvature (Guha et al., 2019). A common mechanism of permeabilization involves oligomerization on membranes, which may be the phenomenon we observed with BefA puncta that form on eukaryotic cells. Alternatively, these BefA puncta may represent endocytic or exocytic responses of cells repairing BefA-induced membrane damage (Brito et al., 2019).

Lipid membrane repair is critical for the survival of all cells. Animal cells, which lack defensive cell walls, have evolved extensive mechanisms for monitoring and maintaining their membrane integrity, including responding to increases in intracellular calcium and extracellular ATP that result from plasma membrane ruptures (Horn and Jaiswal, 2018). Pancreatic β-cells rely on these same cues to regulate their insulin secretion and thus may be especially sensitive to perturbations in their membrane integrity (Noguchi and Huising, 2019). Membrane attack can come from microbial products, such as pore forming toxins and antimicrobial peptides, or from host defense proteins like the pore-forming Reg family antimicrobial lectins. The functions and cellular targets of these host-derived proteins are not fully understood. For example, while Reg proteins are highly expressed and secreted at the apical border of intestinal epithelial cells, where their antimicrobial activity plays a key role in the maintenance of a healthy barrier against microbes (Vaishnava et al., 2011; Wang et al., 2016), they are also expressed in tissues lacking a resident microbial population such as the developing mammalian pancreas (Bartoli et al., 1998; Mally et al., 1994; Perfetti et al., 1996). Reg proteins are also upregulated during β-cell regeneration (Terazono et al., 1988), and are capable of stimulating β-cell expansion (Unno et al., 2002). Type I, or RegI, proteins are associated with β-cell turnover and protection from diabetes in animal models (Okamoto, 1999). In the T1D NOD mouse model, antibiotic-induced disease acceleration was associated with decreased Reg3 expression in both the ileum and pancreas, and upregulated by early life exposure to a protective microbiota (Zhang et al., 2021). Furthermore, RegI and Reg3 proteins are upregulated in humans with diabetes (Astorri et al., 2010; Bacon et al., 2012; Yang et al., 2015a) and are hypothesized to play a role in promoting regeneration and recovery of β-cells during type 2 diabetes remission (Sala et al., 2017). Similarly, another antimicrobial peptide, Cathelicidin, is expressed in rat and human β-cells and treatment with this pore-forming peptide stimulates short term insulin secretion and longer term β-cell expansion (Pound et al., 2015). Until now, neither the pore forming activities of Reg nor Cathelicidin have been linked to their roles in the pancreas. Our insights into the biochemical properties of the bacterial BefA protein demonstrate that diverse membrane-permeabilizing proteins can stimulate β-cell development, pointing to membrane manipulations as a new approach for stimulating β-cell renewal to treat diabetes and motivating future experiments to test whether BefA can prevent or ameliorate diabetes in animal disease models.

Membrane permeabilizing activity is a ubiquitous feature of multispecies microbial communities, representing what we call a Microbial Associated Competitive Activity (MACA) (Wiles and Guillemin, 2020). Membrane disrupting MACAs are detected by host immune sensors such as the NLRP3 inflammasome (Swanson et al., 2019), similar to host immune reception of Microbial Associated Molecular Patterns (MAMPs). We hypothesize that manipulation of membrane integrity is an underexplored mechanism in tissue development and plasticity. The earliest eukaryotic cells, evolving among MACA-rich bacterial communities, would have required membrane defense and counterattack strategies. Extant eukaryotes have elaborate repertoires of MACA mimics, including many membrane permeabilizing antimicrobial proteins (AMPs). Strikingly, these host-derived AMPs, like Reg proteins and Cathelicidin, have a breadth of immunomodulatory activities at lower concentrations and tissue locations distant from where they function in microbial defense (Hancock et al., 2016). We hypothesize that microbiota induction of host defense proteins, along with microbiota-derived secreted proteins, contribute to programs of tissue development, such as early life β-cell expansion.

The developmental plasticity of the pancreas in response to microbial membrane permeabilizing activities may confer a selective advantage by allowing developing animals to match their metabolic capacities to their nutritional environment. Bacterial abundance is a signature of nutrient availability. Membrane permeabilizing activity specifically is a characteristic of bacteria that compete successfully in dense, multispecies communities found in animal digestive tracts. For example, of all the bacterial taxa encoding pore-forming MACPF superfamily members, the most abundant representatives are the Firmicutes, Bacteroidetes, Actinobacteria, and Gammaproteobacteria (Moreno-Hagelsieb et al., 2017), the most common residents of vertebrate digestive tracts. Almost all the BefA homologs we found in human-associated bacteria were produced by the Gammaproteobacteria, which are metabolically versatile, fast-growing, “weedy” species that are often abundant members of the neonatal gut microbiome, but become rarer in healthy adults (Bokulich et al., 2016; Yassour et al., 2018). The exposure of developing pancreatic cells to bacterial-derived membrane-perturbing MACAs and to self-produced antimicrobial defense MACA mimics (such as Regs) would provide information about the abundance of bacteria and by extension nutrients in the environment.

The recent surge of type 1 diabetes in high income countries is strongly associated with increased hygiene measures that reduce microbial exposure during early infant life. In a refinement of the hygiene hypothesis (Strachan, 1989), we hypothesize that increased early life hygiene has resulted in mismatched microbial and nutritional developmental cues. Low exposure to membrane-perturbing MACAs would set up developmental programs for conditions of nutrient scarcity, restricting the β-cell population, and leaving individuals ill-prepared for future diets of caloric excess and with limited β-cell reserves to counteract future autoimmune attacks or hyperglycemia that would drive diabetes development. Understanding how different membrane permeabilizing activities stimulate β-cell expansion will spur new strategies to prevent or reverse the consequence of developing with mismatched microbial and nutritional cues.

## Acknowledgements

We thank: B. Nolen for providing us with purified F-actin and cortactin to use in our co-pelleting assays, for which his lab also provided guidance, J. Davidson for assistance with protein purification, S. Hanson for direction in the construction of the mNeonGreen fused BefA constructs, L. Hooper for advice in purification of Reg proteins and antimicrobial assays, T. Wiles for advice in the construction of BefA constructs, C. Tahmessbipour for assistance with IHC of pancreas cross-sections, K. Ost and A. Weis for critical reading of the manuscript, R. Sockol and the University of Oregon Zebrafish Facility for maintenance of zebrafish lines, J. Williamson and the University of Utah Animal Facilities, for the maintenance of both GF and SPF mice, M. Bridge for training and help with use of the AxioScan slide scanning microscope at the University of Utah Cell Imaging Core, The Berkeley Center for Structural Biology, which is supported in part by the National Institutes of Health, National Institute of General Medical Sciences, and the Howard Hughes Medical Institute, the Advanced Light Source, which is supported by the Director, Office of Science, Office of Basic Energy Sciences, of the U.S. Department of Energy under Contract No. DE-AC02-05CH11231, the Pilatus detector, which was funded under NIH grant S10OD021832, and the ALS-ENABLE beamlines, which are supported in part by the National Institutes of Health, National Institute of General Medical Sciences, grant P30 GM124169.

## Author Contributions

Conceptualization, J.H.H., M.S.M, E.G.S., K.G.; Methodology, J.H.H, M.S.M., E.G.S., E.S.W., P.J., R.P., J.R., K.G.; Software, P.J, R.P.; Analysis, J.H.H., M.S.M., E.G.S., E.S.W., P.J., K.K., S.J.R; Investigation, J.H.H., M.S.M., E.G.S., E.S.W., P.J., K.K., D.D., R.B., S.J.R.; Resources, P.J., R.P., R.B., S.J.R; Writing – Original Draft, J.H.H., M.S.M., E.G.S., E.S.W., K.G.; Writing – Review & Editing, J.H.H., M.S.M., E.G.S., E.S.W., P.J., L.C.M., R.P., S.J.R., J.R., K.G.; Supervision, R.P, L.C.M., J.R., K.G.

## Methods

### Protein Biochemistry

#### BefA protein expression & purification

The *befA* gene was expressed and purified as previously described (Hill et al., 2016). Additionally, BefA protein containing selenium methionine was produced as described by Van Duyne et al. and purified using the same methods as native BefA (Hill et al., 2016; Van Duyne et al., 1993). The nucleotide sequences corresponding to the Befa^99^ and BefA^133^ truncations were amplified using the same reverse PCR primer previously published for amplifying BefA (Hill et al., 2016), which was paired with the following forward PCR primers for each truncation respectively: 5’-GGCCATATGATGaagacggcgaaagaggcgagg-3’ and 5’-GGCCATATGATGggttatgcggtgttcgattcgcgc-3’. A SOE (splicing by overlap extension) PCR reaction (Horton et al., 1990) was used to fuse the *mCherry* or *mneongreen* genes to the 5’ end of either the *befA* or the *befA^99^* gene with a six amino acid linker (AAAGGG) connecting them. Each construct was then cloned into the pET-21b+ plasmid (Novagen, Darmstadt, Germany) and expressed and purified using a C-terminal HisTag^®^ as previously described for the native BefA protein (Hill et al., 2016).

#### Human Reg3α protein expression & purification

Human Reg3α (hReg3α) lacking the N-terminal inhibitory pro-peptide was expressed and purified from a pET3a vector (Addgene plasmid ID 64937, deposited by the laboratory of Dr. Lora Hooper) in *E. coli* BL21 DE3. hReg3α was purified based on the denaturing protocol in (Cash et al., 2006b), with some differences provided by personal communication with the Hooper laboratory. Briefly, hReg3α was expressed in a 1 L culture of *E. coli* with 1mM IPTG for 3 hrs at 37°C shaking, cells were pelleted and stored at -20°C. The cell pellet was resuspended at 4°C in 50 mL of 20 mM Tris pH 7.5, 10 μM EDTA pH 8 and 1% Triton x-100. All subsequent steps were performed at 4°C unless otherwise noted. Cells were lysed using a sonicator, pelleted, supernatant removed and the pellet resuspended using a dounce homogenizer in 100 mL of 20 mM Tris pH 7.5, 10 μM EDTA pH 8, 500 mM NaCl and 1% Triton x-100. The pelleting step was repeated, the supernatant was removed and resuspend again in the previous solution. The solution was centrifuged and the pelleted inclusion bodies were resuspended gently in 20 mL of 7M guanidine-HCl, 10 μM reduced glutathione, 20 mM Tris pH 8.0 and 2 mM EDTA. The solution was rotated for 24 hrs at room temperature. The solubilized inclusion bodies were spun down and 4 mL of the supernatant was added, dropwise, to 200 mL of refolding buffer: 50 mM Tris pH 8.0, 10 mM KCl, 2 mM MgCl_2_, 2 mM CaCl_2,_ 240 mM NaCl, 500 mM Guanidine-HCl, 400 mM sucrose, 500 mM arginine-HCl, 1 mM reduced glutathione, 0.75% Triton x-100 and 0.1 mM oxidized glutathione. Solution was left to stand, covered for 24 hrs. The solution was centrifuged and the supernatant collected and then a series of dialysis steps were performed. First, the protein solution was dialyzed against 25 mM Tris pH 7, 2 mM CaCl_2_ and 25 mM NaCl for 8-24 hrs, then repeated with a fresh batch of dialysis solution. Then the solution was dialyzed into 25 mM MES pH 6.0, 2 mM CaCl_2_, and 25 mM NaCl for 8-24 hrs in preparation for cation exchange chromatography. After dialysis finished, the solution was centrifuged and the supernatant collected. The solution was passed over a 2 mL SP-Sepharose column (cation exchange) equilibrated with the last dialysis solution. The column was washed with 25 mL of 25 mM MES pH 6.0, 2 mM CaCl_2_, 25 mM NaCl and 150 mM NaCl. The protein was eluted with 15 mL of 25 mM MES pH 6.0, 2 mM CaCl_2_, 25 mM NaCl and 400 mM NaCl. The eluted protein was dialyzed into water or 25 mM MES pH 5.5 and 25 mM NaCl and concentrated using a Vivaspin 20 centrifugal concentrator with a 3 kDa molecular weight cutoff (Millipore Sigma, St. Louis, MO).

#### BefA crystallization

Purified BefA and selenium methionine substituted BefA (SeMetBefA) were crystalized from a starting concentration of 10-16 mg/mL in ddH_2_0 in a reservoir solution of: 24-25% PEG 3350 (Hampton Research, Aliso Viejo, CA), 0.1M citric acid pH 3.5, and for semetBefA, 1mM TCEP (tris(2-carboxyethyl)phosphine). Hanging drop vapor diffusion with ratios of protein to reservoir solution of either 1:1 or 2:1, resulted in crystals within 7-10 days. Crystals were cryoprotected in the reservoir solution plus 20% PEG 200 (Hampton Research, Aliso Viejo, CA) and were flash frozen in liquid nitrogen for data collection at the Advanced Light Source in Berkeley, CA beamline 5.0.2 using the Pilatus detector at a wavelength of 1 Å.

#### BefA structure determination

The space group is C2 with a=94.49, b=64.28, c=42.60 Å and β=113.43, with one molecule in the asymmetric unit. Various diffraction data sets were collected to ∼1.3 Å resolution at the Advanced Light Source in Berkeley, CA beamline 5.0.2 using the Pilatus detector at a wavelength of 0.9795 Å and were integrated and scaled using the program package HKL2000 (Otwinowski and Minor, 1997). The initial structure determination was based on a single selenomethionine data set using Autosol from the PHENIX program package (Liebschner et al., 2019) using the SIRAS approach. The Autosol/Autobuild procedures produced a model with 202 residues identified and placed in 5 fragments, including 203 water molecules. Rwork/Rfree for that model were 0.198/0.210 at 2.0 Å resolution. Several cycles of manual model corrections using the program Coot (Emsley et al., 2010) against the SeMet data set resulted in a nearly complete model (residues 36-258) with Rwork/Rfree 0.174/0.197 at 1.31 Å resolution (Table S1).

#### BefA Structure Refinement

Initial rigid body refinement using the semetBefA structure, merged data from two native data sets and Refmac 5 (Collaborative Computational Project, 1994) resulted in Rwork/Rfree values of 0.297/0.298. Manual model building was performed using Coot 0.8.1 (Emsley and Cowtan, 2004) and the bulk of the refinement was carried out using PHENIX 1.9-1692 (Adams et al., 2010). The high-resolution cutoff was determined by the method of Karplus and Diederichs (Karplus and Diederichs, 2012) using CC1/2 of > 0.3 and completeness of > 50% in the highest resolution shell. Two citrate molecules, found in the crystallization conditions at 100 mM, are bound near Ser218 (Figure S5). They likely coordinate a metal that presents as a positive electron density peak if left out. Attempts to place a magnesium ion between the two citrates, as our best guess, resulted in higher Rwork/Rfree values, and therefore was left out. In addition, the distances between the metal and the citrate oxygens are between 1.4-1.7 Å, which is particularly short. In later stages of refinement, hydrogens, and individual anisotropic B-factors were included and all decreased the Rwork/Rfree values. The final refinement round resulted in Rwork/Rfree of 0.132/0.154 and is included in Table S1. The structure is deposited in the PDB as 7RFQ.

#### Thermofluor melting temperature assay

The thermofluor assay was performed as previously described (Henderson et al., 2013) with the following alterations. A final concentration of 0.3-0.5 mg/mL of BefA or lysozyme (chicken egg white lysozyme, VWR, Radnor, PA) was incubated on ice with 16X SYPRO Orange protein dye (Thermo Fisher Scientific, Waltham, MA), 150 mM NaCl, and 25 mM HEPES or Tris (pH 7-7.5). 20 μL samples were pipetted into a 364 well PCR plate and covered with a clear, optically transparent seal for reading fluorescence. Melting temperature data for BefA and lysozyme, as a control, were collected on a StepOnePlus real time PCR machine (Thermo Fisher Scientific, Waltham, MA). Reactions were run using the following thermal denaturation protocol: 4°C for 1 min, slow ramp rate (∼0.03 °C/sec) from 4°C to 80°C with fluorescence data collected ∼8.5 sec. Fluorescence data was plotted against temperature and approximate melting temperature (T_m_) determined using the following equation: max – ½*(max – min). Max refers to fluorescence intensity maximum value (peak) and min refers to minimum intensity value prior to intensity increase. The BefA thermofluor assay was run on two separate days with a total of 12 replicates. The lysozyme thermofluor assay was run in triplicate on two separate days.

#### Actin co-pelleting assay

The actin co-pelleting assay was performed as described (*10*) with the following changes: BefA, BefA^99^, and cortactin (gift from Dr. Brad Nolen), were pre-clarified to remove aggregate proteins by spinning at 65,000 rpm for 30 min at 4°C in a TLA100 rotor (Beckman Coulter, Brea, CA) before incubating with F-actin. F-actin filament formation was encouraged by incubating purified rabbit actin (gift from Dr. Brad Nolen) in a solution of 10 mM imidazole (pH 7.0), 50 mM KCl, 1 mM EGTA, 1 mM MgCl_2_ and 1 mM DTT at room temperature for 45 min. The supernatants of the clarified proteins were incubated at a final concentration of 2.5 µM, with or without F-actin at 5 µM, in the buffer described above for 1.5 h at room temperature. All reactions were centrifuged at 65,000 rpm for 30 min at 4°C in a TLA100 rotor. The supernatant (unbound) and pellet (bound) proteins were separated and analyzed by SDS-PAGE and stained with coomassie brilliant blue.

#### PIP strip Assay

BefA, negative control AimA (Rolig et al., 2018) and positive control PI(4,5)P_2_ Grip (Echelon catalog number G4501) all contain 6X-His tags and were incubated with individual membrane strips or arrays at a final concentration of 0.75 μg/mL for 1 hr at RT as directed by the manufacturer (Echelon Biosciences, Inc., Salt Lake City, UT). For visualization, an anti-His rabbit polyclonal antibody (ABM, Richland, BC, Canada) in combination with a polyclonal goat anti-rabbit HRP conjugated secondary antibody (Thermo Fisher Scientific, Waltham, MA) were used to detect the presence of His-tagged proteins after washing the membranes with 1X PBS-T (phosphate buffered saline with 0.1% Tween-20) as directed. Membrane and PIP lipid strips were performed with BefA 3-5 times, once with AimA and once with Grip for the PIP lipid strip. The membrane lipid arrays with or without calcium were performed once with BefA.

#### Vesicle preparation, treatment, and quantification

Vesicles were made as previously described using electroformation (Angelova M.I., 1992; Veatch, 2007), with the following lipid molar ratio to obtain red fluorescent vesicles: 99.5 % 1,2-dioleoyl-sn-glycero-3-phosphocholine (DOPC) (Avanti Polar Lipids Inc., Alabaster, Alabama) and 0.5 % Texas Red 1,2 Dihexadecanoyl-sn-Glycero-3-Phosphoethanolamin (Texas Red DHPE) (Thermo Fisher Scientific, Waltham, MA). Fresh vesicles in 0.1 M sucrose were made the previous day and incubated with either the same volume of water (control) or BefA in water to a final concentration of 2 µM for 35 min at room temperature. Treated vesicles were loaded into a 350 µL glass cuvette (Starna Cells, part number 3-3.45-SOG-3, Atascadero, CA) and imaged on a home-built light sheet fluorescence microscope described in Jemielita et al. 2014 (Jemielita et al., 2014). Four, three dimensional stacks of fluorescent images, spaced 1 µm apart, were collected across the middle of the cuvette 35 and 50 min after addition of BefA protein or water (control). Phenotypes, such as percent of multi-vesicular vesicles and vesicle concentration, were quantified using a custom-made Python script (available here: https://github.com/rplab/vesicle_detector) that identified potential vesicles and then sorted by hand, removing false positives and sorting features as single vesicles or multi-vesiculated vesicles. Percentages were calculated as the number of multi-vesiculated vesicles divided by the total number of features (sum of multi-vesicular vesicles and single vesicles). The total number of single and multi-vesicular vesicles (three or more vesicles attached) were quantified and graphed in GraphPad Prism.

#### Vesicle dye release assay

The vesicle dye release assay was done according to Jimah et al. with some slight changes (Jimah et al., 2017). Briefly, a total of 1 mg of DOPC lipids (Avanti Polar Lipids Inc., Alabaster, Alabama) in chloroform was dried under vacuum for 3 hrs, resuspended in 1 mL of 20 mM carboxyfluorescein, 25 mM Tris pH 7.5 and 25 mM NaCl with a brief vortex, sealed with parafilm and left overnight at 4 °C. The next day, lipids were extruded 10 times through 100 nm or 400 nm pore size membranes using a LIPEX 1.5 mL Extruder (Transferra Nanosciences Inc., Burnaby, B.C., Canada) at 50 °C to form vesicles of a more uniform size. Extruded vesicles with dye inside were separated from free dye by using a PD-10 column. Samples were loaded into a black 96 well, flat bottom plate with a clear bottom (Corning, Corning, New York). Samples were read using a BMG Labtech FLUOstar Omega plate reader (fluorescence ex/em 485/520 nm). Fluorescent measurements of vesicles without treatment were read initially, treatments were added, and then read for 1 hr every 2 min. Finally, 1% Triton x-100 was added to obtain maximum fluorescence values. The initial readings were subtracted from each curve and the percent of max was calculated by taking each treatment reading and dividing it by the final, maximum Triton x-100 value. Data was plotted in GraphPad Prism using mean and standard deviation.

#### CyQUANT assay for bacterial membrane stability

Bacterial strains were maintained in 25% glycerol at −80 °C. Bacteria were directly inoculated into 5 mL lysogeny broth (LB) media (10 g/liter NaCl, 5 g/L yeast extract, 12 g/L tryptone, 1 g/L glucose) and grown for ∼16 hrs (overnight) with shaking at 30 °C. Bacterial overnight cultures were subcultured and grown to an approximate optical density at 600 nm of 0.1 to 0.3. The bacterial cells were washed twice by centrifugation, aspiration of supernatant, and suspension in zebrafish embryo media (pH=5.5); after the second aspiration step, bacterial cells were concentrated by suspending the pellet in a volume of sterile embryo media (pH=5.5) that was a quarter of the starting volume. CyQUANT Direct Red (Invitrogen; product no. C35014) 2X Reagent was prepared fresh as directed by the kit’s protocol by diluting CyQUANT™ Direct Red nucleic acid stain (1:250) and CyQUANT Direct Red background suppressor (1:50) in embryo media (pH=5.5). In a sterile 96-well tissue culture-treated black flat-bottom microplate (Greiner Bio-One; product no. 655090), appropriate volumes of bacteria, or sterile embryo media (pH=5.5) in controls, were mixed in triplicate with 50 μL/well of CyQUANT 2X Reagent. Cells and CyQUANT 2X Reagent were incubated in the dark at 30°C for one hour. A FLUOstar Omega microplate reader (BMG Labtech, Offenburg, Germany) was used to obtain baseline fluorescence measurements with 584 nm excitation and 650 nm emission filters before addition of treatment solutions. Treatments were added to each well to achieve desired final concentrations in a total volume of 100 μL/well. Final concentrations of treatments were as follows: 7 μM BefA, 3 μM Reg3α, and 0.08% SDS. Fluorescence measurements were then taken every 10 min for 400 min at 30°C, the plate was shaken briefly before each reading. Unfortunately, addition of BefA^99^ in this assay caused dye precipitation which confounded our analysis. We also tried using BefA in a propidium iodide (PI) assay, another dye often used for assessing membrane permeability, and these efforts were also inconclusive due to natural autofluorescent in the presence of propidium iodide.

#### Bacterial imaging and quantification

*Bacillus* sp. (strain index # ZOR0058) a contaminant isolated from a GF zebrafish flask, was grown overnight at 30°C, shaking, from a glycerol stock. The culture was back-diluted in the morning (approximately 16 hrs later) at a 1:50 dilution and grown at 30°C shaking for approximately 1.5 hrs. 1 mL of bacterial culture was pelleted and gently washed into 500 μL of PBS. Purified mCherry-BefA or mCherry control protein was incubated with bacterial cells at 1 μM for 30 min at room temperature. SYBR Green nucleic acid stain (Thermo Fisher Scientific, Waltham, MA) was added to cells at 1X final concentration in PBS and incubated at room temperature for 20 min. 100 μL of cell mixture was loaded into multi-well microscope slides with a loose lid. Images in three channels, DIC, SYBR Green (ex/em 475/525 nm), and mCherry (ex/em 575/625 nm) were acquired using a GE DeltaVision Ultra fluorescent microscope using a 60x magnification, 1.42 numerical aperture, Plan Apo N, oil immersion objective. The DIC channel was taken at 50 msec exposure with 10% power, the SYBR Green channel was taken at 100 msec exposure with 10% power and the mCherry channel was taken at 75 msec exposure with 32% power. 10-15 original images (1024 x 1024 px) were taken of each sample and processed identically within each channel using Fiji (ImageJ). Each channel was pseudocolored using the channels tool (DIC, gray; SYBR Green DNA, green; mCherry proteins, magenta) and the min and max displayed values set for the first two channels (DIC min/max 702/3934 and green 575/15605), while magenta/protein channel was background subtracted using a rolling ball radius of 50 px. A 20 x 20 μm square was cropped around representative cells, and then each image scaled identically in Adobe Illustrator CC 2018.

For quantification of BefA binding, all cell regions in the two treatments that were not overlapping with other cells were quantified for BefA binding using Fiji. In the DIC channel, the segmented line tool at 35 px wide was used to draw along the length of each cell, clicking at each segment. For every image, a line of roughly the same length as the average bacterial cell was drawn to capture the background intensity. The mean gray value and standard deviation was measured for each line using the Measure analysis tool. The mean background values were subtracted from each value taken along a cell and were plotted in GraphPad Prism.

#### Anti-BefA antibody preparation and testing

Two rabbit polyclonal antibodies were produced through Cocalico Biologicals (Stevens, Pennsylvania) using their standard 56-day protocol (three boosts), ending in exsanguination. Briefly, BefA with 6x His affinity tag was purified as previously described (Hill et al., 2016), separated from impurities by SDS PAGE and the BefA band excised. The gel slices containing 1-2 mg protein were mailed to Cocalico Biologicals. Two rabbits were chosen based on pre-bleeds showing little response to other antigens of similar molecular weight to BefA on western blots. The rabbits were injected with BefA antigen and two bleeds (∼6-8 mL each) plus the final bleed (∼30-50 mL each) were collected. The bleeds were analyzed via western blot for reactivity against purified BefA (full-length, truncated BefA99 and fluorescently-tagged forms) as well as endogenous BefA from bacterial cultures. 1 μg of purified protein or 25 μL of bacterial culture was loaded on an SDS PAGE gel and transferred to a PVDF membrane. The membrane was blocked in 5% milk in Tris-Base Saline Tween-20 (TBST) and then probed with our antibody bleeds against BefA at a 1:500 concentration in 5% milk/TBST overnight at 4°C. To visualize the resulting BefA bands, a secondary Anti-rabbit IgG HRP-linked antibody (Thermo Fisher Scientific, Waltham, MA) was used at 1:1000 for 1 hour at room temperature. Additional samples of bacteria cultures that lacked endogenous BefA were run simultaneously with our other samples to confirm that the antibody is specific.

### Animals

#### Gnotobiotic zebrafish husbandry and protein treatment

All zebrafish experiments were performed using protocols approved by the University of Oregon Institutional Care and Use Committee (protocol number 14-14RR) and followed standard protocols. Wild-type (ABC x Tu strain), *Tg(−1.0insulin:eGFP)* (referred to as *ins:gfp*) (diIorio et al., 2002), *Tg(ins:mCherry)jh2* (Pisarath et al., 2007), *myd88^-/-^ (31*), and *sox9bfh313* (referred to as *sox9b^-/-^*) (*35*), npas4l (referred to as *cloche) (38, 39),* and *Tg(nxk2.2:eGFP)* (Pauls et al., 2007) zebrafish embryos were either raised conventionally or derived GF as previously described (Bates et al., 2006; Hill et al., 2016). Following GF derivation, fish were mono-associated with bacterial strains as previously described (Bates et al., 2006; Hill et al., 2016) or treated with a given purified protein. For experiments involving the treatment of larvae with sterile purified protein, the protein was either added directly to the embryo media at a final concentration of 500 ng/mL at 4 dpf, unless specified otherwise, or delivered using oral microgavage at the concentration indicated in the text with rigorous sterile technique. Larvae were incubated with the protein of interest for 48 hrs or until 6 dpf.

#### Quantification of zebrafish β-cells

Larvae were fixed in 4% PFA, antibody stained, and imaged using a confocal microscope as described previously (Hill et al., 2016) to quantify total β-cell numbers. For experiments utilizing *Tg(−1.0insulin:eGFP)* larvae, antibody staining was conducted as previously described (Hill et al., 2016). For experiments utilizing *myd88^-/-^, sox9b^-/-^*, and *cloche* larvae, they were first permeabilized using an acetone freeze step for 7 min at -20° C before staining with guinea-pig anti-insulin (Dako, Carpinteria, CA), Alexa Fluor 488 or 647 goat anti-guinea-pig (Thermo Fisher Scientific, Waltham, MA), and either TO-PRO-3-Iodide (Thermo Fisher Scientific, Waltham, MA) or DAPI (Thermo Fisher Scientific, Waltham, MA).

#### Larval zebrafish oral microgavage and cardiac valley injections

For oral microgavage, 4 or 6 dpf larval zebrafish were anesthetized, mounted in sterile 4% methylcellulose and gavaged as described previously (Cocchiaro and Rawls, 2013). Proteins at the concentrations specified in the text were administered to each larval zebrafish with a 4.6 nL volume at an injection rate of 23 nL/sec using a Nanoject II Auto-Nanoliter Injector (Drummond Scientific Company, Broomall, PA). To visualize the global localization of BefA *in vivo*, mCh-BefA, mCh, mNG-BefA, or mNG were administered at a concentration of 1 mg/mL. All images were taken no longer than 2 hrs after gavage using the wide-field feature of a LEICA DM6 confocal microscope (Leica Microsystems Inc., Buffalo Grove, IL).

For cardiac valley injections, 4 dpf larval zebrafish were anesthetized and injected directly into the cardiac valley with 2 pg of BefA or negative control vehicle again using the Nanoject II Auto-Nanoliter Injector (Drummond Scientific Company, Broomall, PA) as previously described (Wiles et al., 2009). Zebrafish were revived and housed in separate wells of a 24-well plate after the injection to monitor health. To ensure that gavaged or injected GF larvae were not contaminated with microbes during these procedures, a sample of embryo media and homogenized gastrointestinal tracts from some larvae were plated onto tryptic soy agar (TSA) or luria broth (LB) agar at the time of harvest, typically 48 hrs later.

#### BODIPY feeding and analysis

A BODIPY/egg yolk mixture was prepared by emulsifying sterile 5% chicken egg yolk in zebrafish embryo media via sonication for 3-5 min (Carten et al., 2011). BODIPY dissolved in DMSO (Thermo Fisher, Waltham, MA) was quickly added to the emulsified egg mixture at a final concentration of 6.4 µM and vigorously vortexed for 3-5 min. 1ml of the BODIPY/egg yolk mixture was added to 4dpf zebrafish in a sterile 12-well plate (10 zebrafish/well) for 5 hrs. Zebrafish were covered in foil during their exposure to the BODIPY/egg yolk mixture. To visualize the patency of the pancreatic ductal system, the confocal feature of a LEICA DM6 confocal microscope (Leica Microsystems Inc., Buffalo Grove, IL) was used for imaging live zebrafish administered BODIPY/egg yolk. With DIC, the pancreas was located and z-stack images throughout the entire head of the pancreas were acquired. The presence or absence of BODIPY within the pancreas was assessed by compiling the z-stacks taken of the pancreas into maximum projections.

#### Gnotobiotic mouse husbandry

SPF C57BL/6J and Swiss Webster mice were originally purchased from Jackson Labs and reared in the SPF facility at the University of Utah. Germfree mice were reared in sterile gnotobiotic chambers in the germfree facility at the University of Utah. Microbial sterility was determined every 3 weeks and at sacrifice by plating and PCR. For *E. coli* Nissle mono-associations, breeder pairs were inoculated via oral gavage with approximately 10^6^ CFU of a given strain. Mono-associations for each strain were performed in separate gnotobiotic chambers to prevent cross-contamination. Successful and continual Nissle colonization was confirmed by periodically plating feces to monitor the presence of a single colony morphology, and was further confirmed by 16S sequencing. Contamination was also ruled out in the same manner. Periodic PCR amplification of the *befA* gene was also used to monitor each mono-association experiment over time. All experiments were performed in accordance with federal regulations as well as the guidelines for animal use set forth by the University of Utah Institutional Animal Care and Use Committee under protocol numbers 17-04009 & 20-03006.

Oral gavage delivery of BefA to neonates was performed three times at P3, 5, and 7, as described (Francis et al., 2019). BefA was delivered at a final concentration of 500 μg/mL. The same concentration was also used to deliver BefA via neonatal IP injection at P3, 5 and 7. C57B6 mice were used for oral gavage experiments, and assayed at P12 and Swiss Webster mice were used for IP injection experiments and assayed at P21.

#### Quantification of mouse beta-cells and alpha-cells

Entire wild-type litters of either C57BL/6J or Swiss Webster of the indicated gnotobiology were collected at neonatal or adult ages as indicated in the text, and immediately euthanized before harvesting the pancreas. Freshly dissected pancreata were weighed before fixing overnight in 10% buffered zinc formalin or Z-FIX (Anatech LTD, Battle Creek, MI) at RT. Fixed pancreata were rinsed 3 times in PBS before dehydrating in increasing concentrations of EtOH. Dehydrated samples were paraffin embedded and for each sample at least 3 longitudinal sections of 6 um thickness and 100 um apart (for neonates) or 200 um apart (for adults) were cut and mounted onto glass slides for analysis. Slides were then deparaffinized, rehydrated, permeabilized, blocked in PBS with 5% goat serum (Jackson ImmunoResearch Laboratories, INC., West Grove, PA) at RT for one hour, and stained with a guinea pig anti-insulin antibody (Dako, Carpinteria, CA) at a final concentration of 1:200 overnight (ON) at 4°C. Primary antibody was washed in PBS with 0.1% Tween 20 (PBT) before applying a horseradish peroxidase (HRP) conjugated anti-guinea pig antibody (Jackson ImmunoResearch Laboratories, INC., West Grove, PA) at a final concentration of 1:250 for 2 hrs at RT or ON at at 4°C. After rinsing with PBT, HRP was detected using a DAB substrate kit (Abcam, Cambridge, MA). Counterstaining of nuclei was performed using Meyer’s Hematoxylin (Millipore Sigma, St. Louis, MO). Slides were dehydrated before mounting with Permount (Fisher Scientific, Waltham, MA). An Axio Scan.Z1 slide scanner (Zeiss, White Plains, NY) brightfield microscope was used to image and subsequently stitch entire stained pancreatic cross sections into single images for analysis. The open access software Fiji was used to quantify both insulin-positive area and total cross-sectional area for each section. The ratio of insulin area:total pancreatic area was determined for each section and then averaged across three sections for each mouse. For adult mouse beta-cell quantifications, the same procedure was used, except that mice were harvested at 6-8 weeks of age and beta-cell mass was calculated by multiplying the final ratio of insulin area:total pancreatic area by pancreas weight.

For quantifying alpha-cell mass, immunofluorescence staining, also with the same guinea pig anti-insulin antibody, as well as a mouse anti-glucagon antibody (1:500, Sigma Aldrich, St. Louis, MO) to visualize beta-cell and alpha-cell content. Alexa-flour 647 goat anti-guinea pig (1:500, Thermo Fisher Scientific, Waltham, MA) and Alexa-flour 488 goat anti-mouse (1:500, Thermo Fisher Scientific, Waltham, MA) secondary antibodies were applied. Sections were counterstained with DAPI (1:1500, Thermo Fisher Scientific, Waltham, MA) and mounted for imaging in Prolong Diamond Antifade Mountant (Thermo Fisher Scientific, Waltham, MA). Total insulin and glucagon positive area was calculated for each section using FIJI (Schindelin et al., 2012). The total area covered by pancreatic tissue was determined from DAPI staining, allowing for the percentage of α-cell and β-cell area to be calculated for each section. The ratio of glucagon positive area to either whole pancreatic area or insulin positive area for each cell type was determined in at least 3 cross sections for each mouse by averaging across each section.

#### Quantification of blood glucose and insulin levels from mice

Blood glucose readings were taken in the morning from nonfasted mice at the same time across treatment groups from tail vein samples using a Contour Next EZ glucometer and Contour Next Blood Glucose Test Strips. Serum insulin (also collected from tail vein blood) was measured using the Crystal Chem (Elk Grove Village, IL) Insulin ELISA Kit. For each mouse, two separate 5 uL serum samples were assayed and averaged for the final reported value.

### Cell Culture

#### Zebrafish primary islet dissection and treatment

Gut dissections were performed on 4 dpf GF *Tg(−1.0insulin:eGFP)* larval zebrafish as previously described (Bates et al., 2006). Since the pancreas is tightly associated with the gut, it came along readily with dissected gut tissue. The head of the pancreas was dissected away from the rest of the gastrointestinal organs, so all that remained was the primary islet and a small amount of surrounding exocrine tissue. We found that leaving this small amount of supporting tissue was necessary to keep the islet intact for accurate β-cell quantification. Dissections also likely included small amounts of extra-pancreatic tissue, which varied between dissections, but may have included cell types of the following: ducts, gallbladder, and intestine. Each dissection was transferred into a sterile PCR tube containing 50 uL of Leibovitz’s L15 Media with GlutaMax Supplement (L15) (Thermo Fisher Scientific, Waltham, MA). Protein treatments were added at a final concentration of 500 ng/mL and allowed to incubate at room temperature for 48 hrs. Plating to LB agar at the end of the treatment period allowed us to test sterility of the preparation. Any bacterial contamination was also obvious by color change of the pH indicator in the L15 media. Following treatment, single islets were mounted directly onto a slide with 5 µL of ProLong (Molecular Probes, Eugene, OR) and a coverslip. Each islet was imaged immediately after mounting on a confocal microscope and then analyzed as previously described for *in vivo* β-cells.

#### Preparation of primary zebrafish gastrointestinal cell culture

Gut dissections were performed, as described above, on 4-6 dpf GF Wild-type larval zebrafish, except that an effort was made to preserve other digestive tissues such as pancreas and liver. Dissected tissue of 45-60 larval zebrafish (per group/tube) was collected in 1mL Leibovitz’s L15 media (Fisher Scientific, Waltham, MA) with Penicillin/Streptomycin and supplemented with 15% fetal bovine serum (Fisher Scientific, Waltham, MA). Tubes were gently spun down at 300xg for 3 min to remove the supernatant and the pellet was rinsed 2x with 1mL of 1X HBSS (Fisher Scientific, Waltham, MA). To gently dissociate the dissected tissue into individual cells or small cell clusters, 400 ul of 1X TrypLE (Thermo Fisher Scientific, Waltham, MA) warmed to 30℃ was added and mixed with the pellet by pipetting every 5 min for 15 min. To stop the dissociation reaction, 1mL of cold 1% BSA/HBSS was added and the tubes were spun down at 300xg for 3min. The supernatant was removed and the pellet rinsed twice with 1% BSA/HBSS. The pellet of cells was then re-suspended in 200ul of 15% FBS in Leibovitz’s L15 media with Penicillin/Streptomycin and aliquoted directly onto poly-D lysine coated coverslips (Corning, Corning, NY) in 24-well tissue culture plates.

#### BTC-6 Cell culture

βTC-6 cells (*43*) (ATCC, Manassas, VA) were cultured in low glucose DMEM (Dubelco’s Modified Eagle’s Medium) (Fisher Scientific, Waltham, MA) supplemented with 10% fetal bovine serum (FBS) (Fisher Scientific, Waltham, MA). Cells were incubated at 37° C and 5% CO_2_. Media was changed every 24-48 hrs and cells were split every 4-7 days.

Protein treatments were added to the culture media of either βTC-6 or primary zebrafish cells at a final concentration of 0.22 µM. For imaging analysis, cells were split onto poly-D lysine coated coverslips (Corning, Corning, NY) in 24-well plates. Cells were given at least 24 hrs to attach to the coverslips before treating them with mNG-BefA^99^ or mNG for several hrs. Following the incubation, the media was removed and each well gently rinsed twice with sterile PBS to wash away unbound protein before fixing with 4% PFA for 15 min at room temperature. Cells were permeabilized with 0.1% triton in PBS and then blocked in a solution of 3% BSA, 1% DMSO, and 0.01% Triton in PBS for 10 min. Cells were then antibody stained with guinea-pig anti insulin (1:500, Dako, Carpinteria, CA), Alexa Fluor 647 anti guinea-pig (1:1000 Thermo Fisher, Waltham, MA), and DAPI (1:1000, Thermo Fisher Scientific, Waltham, MA). Coverslips were mounted onto a microscope slide with ProLong Diamond mounting media (Molecular Probes, Eugene, OR) and imaged with a LEICA DM6 confocal microscope (Leica Microsystems Inc., Buffalo Grove, IL) to localize mNG. Images taken were randomized across each coverslip.

To localize where BefA bound to βTC-6 cells, cell fractionation followed by Western blotting was used. Either mCh-BefA^99^, or mCh alone, was administered at a final concentration of 0.89 µM and 1.85 µM, respectively, to the βTC-6 cultures and incubated overnight. For analysis, the cell culture media was collected and concentrated to 1 mL. The cells were rinsed 3 times with 1 mL of cold PBS. All the PBS rinses were collected together and concentrated to 500 µL. Lysis buffer containing 0.01% Triton X-100 and protease cocktail inhibitor (Sigma-Aldrich, St. Louis, MO) was added to each culture dish and put into the -20°C freezer for 5 min. After thawing, the cell remnants were scraped from the bottom of the dish and collected in a clean centrifuge tube. Tubes were spun at 14,000 rpm for 15 min to separate the cytosolic components freed upon lysis from the cell debris. The cytosol within the supernatant and the remaining cell debris pellet were collected separately. The protein concentration for all samples (culture media, PBS rinses, cytosol, and pellet) was measured using the Pierce BCA Protein Assay Kit (Thermo Scientific, Rockford IL) and 20 µg of protein from each sample was run on an SDS PAGE gel and transferred to a PVDF membrane. The membrane was blocked in 5% BSA and then probed with a mouse monoclonal antibody to mCherry (Novus Biologicals, Littleton CO) at 1:1000 at 4°C overnight. To visualize the resulting bands, a secondary Anti-mouse IgG HRP-linked antibody was used (Cell Signaling Technology, Dancers MA) at 1:1000 for 1 hr at room temperature.

#### Primary mouse islet collection and culture

Primary islets were isolated from Swiss Webster pups aged P12 as described (Huang and Gu, 2017). Additional purification was done by multiple rounds of hand-picking. Islets were allowed to recover overnight in RPMI supplemented with 10% FBS (Fisher Scientific, Waltham, MA) at 37° C and 5% CO_2_. Before segregating into separate treatment wells for treatments, islets were screened and unhealthy or dying islets were removed. mNG-BefA^99^ and mNG were each added at a final concentration of 250 µg/mL. Glucose was added as a positive control (Vetere and Wagner, 2012) for beta-cell proliferation at a final concentration of 10 µM. Control wells received no additional treatment besides basal RPMI media supplemented with FBS. Islets were incubated for 48 hours, and were supplemented with 10mM EdU (5-ethyl-2’-deoxyuridine) (ThermoFisher Scientific, Rockford, IL) during the final 6 hours of incubation to mark proliferating cells. Islets were fixed and processed as per the Click-iT® EdU Imaging Kit Protocol before mounting in ProLong Diamond Anitfade Mountant with DAPI (ThermoFisher, Rockford, IL). Islets were imaged using an Olympus IX71 microscope and total EdU labeling was quantified.

### Bioinformatics

#### Amino acid alignment of SYLF containing proteins across kingdoms

Amino acid sequences of SYLF containing proteins obtained from NCBI were aligned using Clustal Omega (Goujon et al., 2010; McWilliam et al., 2013; Sievers et al., 2011).

#### Phylogenetic Analysis

The amino acid sequence of the SYLF domain of BefA from *Aeromonas HM21*, starting at residue 99 and continuing to residue 261, was used to screen for homologs across microbial species using NCBI’s BLASTp (Altschul et al., 1997) function. Default search parameters were maintained except to increase the maximum number of aligned sequences displayed to be 20,000, and the Expect Threshold was changed to 1.0. All standard scoring parameters were maintained. Homologs were identified as those with an amino acid sequence identity equal to or greater than 30%, and a query coverage equal to or greater than 80%. For phylogenetic analysis, each bacterial taxa with a putative SYLF domain-containing protein was represented by the hit of highest percent identity to the SYLF domain of *Aeromonas HM21* among isolates within that taxa. Amino acid sequences of putative SYLF domain-containing proteins were aligned by MUSCLE through Genious Prime 2020.1.2 and the predicted SYLF domain from each sequence was used in a subsequent MEGA 7.0 (Kumar et al., 2016) Alignment by MUSCLE (Edgar, 2004). A rooted phylogenetic tree was constructed using Dendroscope 3 (Huson and Scornavacca, 2012). Taxa habitats were manually curated by cross referencing literature.

#### Analysis of SYLF domain-containing protein lengths

A broad search of “SYLF” was performed in NCBI using Geneious Prime 2020.1.2. All Eukarytoic and Prokaryotic sequence files contianing “SYLF” annotations were downloaded. All sequence length data from the broad NCBI search, along with sequence length data from was exported into a comma separated value file for further analysis. To determine the distribution of sizes of SYLF domain containing proteins, a Frequency Distibution analysis was performed using GraphPad Prism version 6.07 for Windows (GraphPad Software, San Diego, California USA, www.graphpad.com) on sequence lengths from Eukaryotes, Prokaryotes, and the BefA homologs using a bin width of 25 amino acids. The frequency of sequences that were equal to or greater than 1000 amino acids were summed for each group (Eukaryotes, Prokaryotes, BefA homologs) and were represented in one bin. The percent of sequences in each bin was determined by dividing the frequency by the total number of sequences within each group. These percentages were then plotted using GraphPad Prism version 6.07 for Windows.

#### Statistical Analysis

Appropriate sample sizes for all experiments were estimated *a priori* using a power of 80% and a significance level of 0.05. From preliminary experiments we estimated variance and effect. For larval β-cell quantification, these parameters suggested using a sample size of 6 in order to detect significant changes between treatment groups. Each of our experiments contained about 10–15 biological replicates or individual fish per treatment group, although some larger experiments had fewer biological replicates due to limited material, making our studies highly powered. Furthermore, entire experiments or technical replicates were repeated multiple times, resulting in pooled data sets of about 20–50 biological replicates. For mouse beta-cell analysis, our power calculations suggested using a minimal sample size of 4, each mouse being one biological replicate, to detect significant changes. Each of our experiments contained at least 5 mice, except for one which was limited to 3 due to extreme technical difficulty. Technical replicates of mouse experiments were carried out 1 to 3 times. These data are represented in the figures as box and whisker plots, which display the data median (line within the box), first and third quartiles (top and bottom of the box), and max and min of the data (whiskers). Plot of mouse data display all biological replicates in the set as dots overlaying the box plot. In box plots of zebrafish data, which contain far more biological replicates, box plot whiskers represent the 95% confidence interval, and any data point falling outside the 95% confidence interval is represented as a solid dot. Pooled data from multiple replicate experiments were analyzed through either the statistical software RStudio (RStudioTeam, 2020) or GraphPad Prism 8. For experiments comparing just two differentially treated populations, a Student’s t-test with equal variance assumptions was used. For experiments measuring a single variable with multiple treatment groups, a single factor ANOVA with post hoc means testing (Tukey) was utilized. Tukey results are displayed as lower-case letters above each treatment group on every box plot. A p-value of less than 0.05 was required to reject the null hypothesis that no difference existed between groups of data. In rare cases, extreme outliers were removed from data sets using the ROUT method for Robust regression and outlier removal developed in GraphPad PRISM (Motulsky and Brown, 2006), using a Q score of 1%.

## Supplemental Information

Supplemental Figures and Legends 1-5

Supplemental Table 1

*Full PBD X-ray Structure Validation Report for BefA*

