## Supplemental Information for "A microbiota membrane disrupter disseminates to the pancreas and increases β-cell mass"

### Figure S1

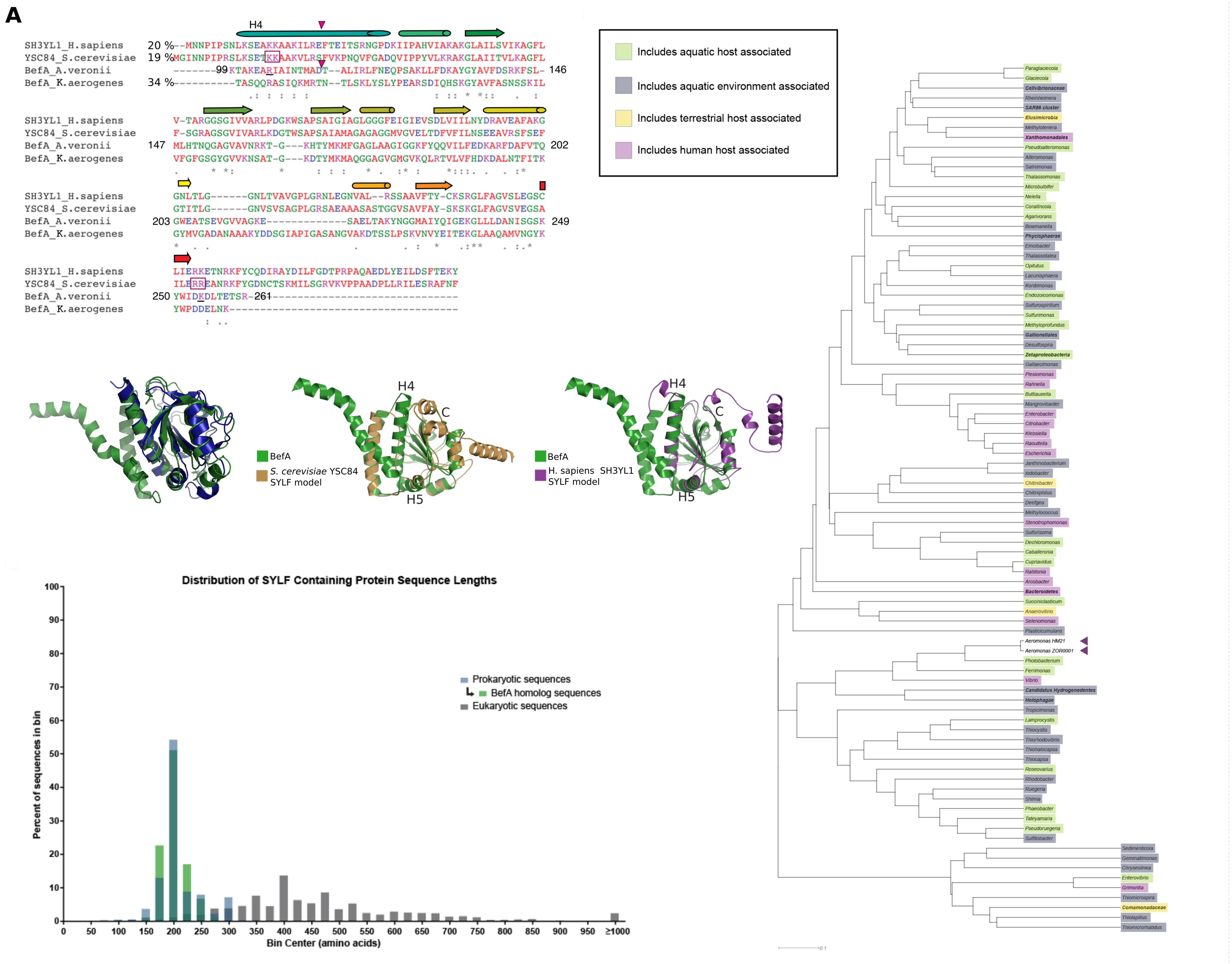
A
H4
SH3YL1_H.sapiens 20 %
YSC84_S.cerevisiae 19 %
BefA_A.veronii
BefA_K.aerogenes 34 %
99
146
147
202
203
249
250
261
Includes aquatic host associated
Includes aquatic environment associated
Includes terrestrial host associated
Includes human host associated
H4
C
H5
BefA
S. cerevisiae YSC84 SYLF model
BefA
H. sapiens SH3YL1 SYLF model
Distribution of SYLF Containing Protein Sequence Lengths
Prokaryotic sequences
BefA homolog sequences
Eukaryotic sequences
Percent of sequences in bin
Bin Center (amino acids)
0
50
100
150
200
250
300
350
400
450
500
550
600
650
700
750
800
850
900
950
≥1000
10
20
30
40
50
60
70
80
90
100
Aeromonas HM21
Aeromonas ZOR0001

### Figure S2

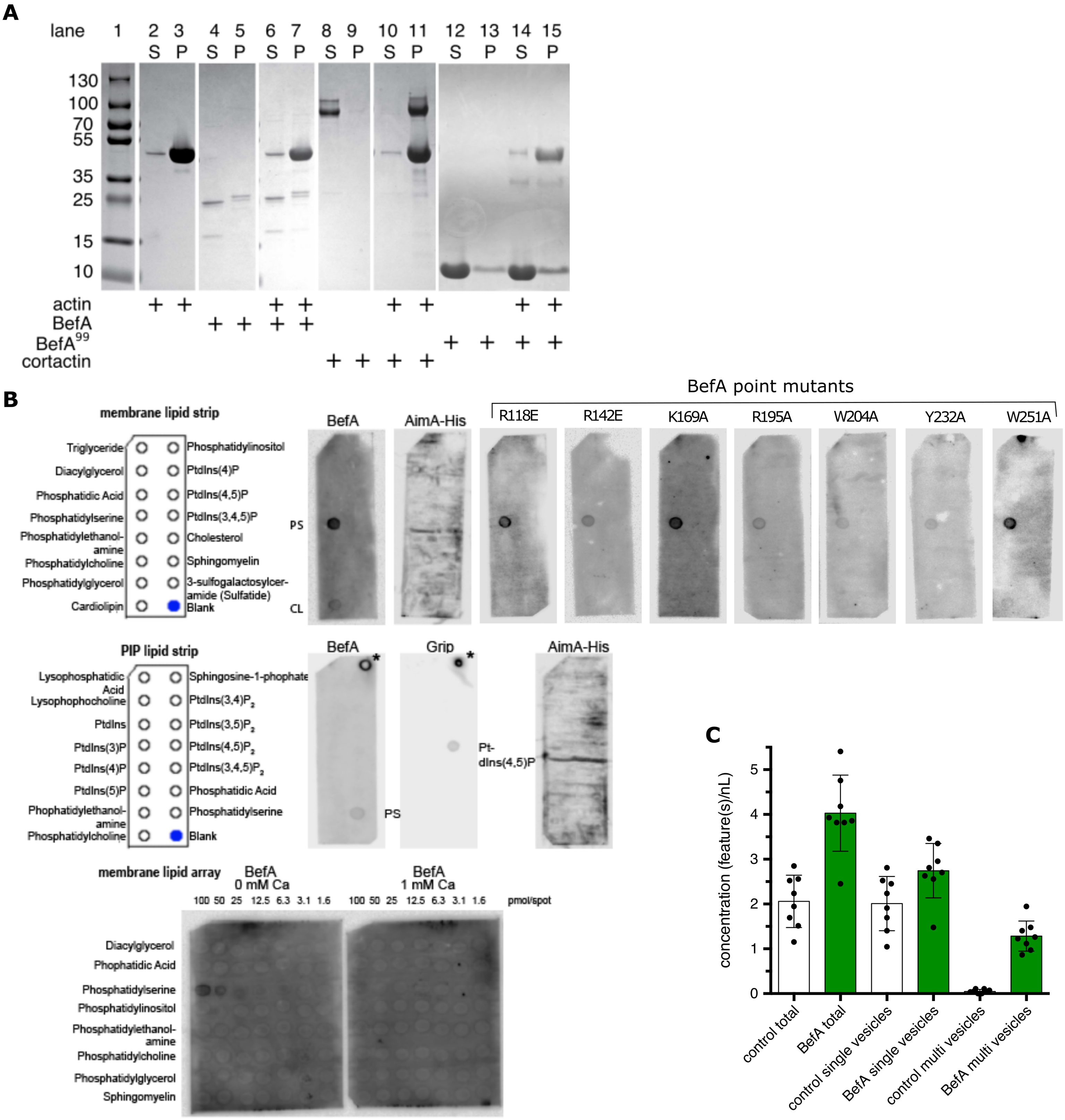

A
lane 1 2 3 4 5 6 7 8 9 10 11 12 13 14 15
S P S P S P S P S P S P S P
130
100
70
55
35
25
15
10
actin
BefA
BefA99
cortactin
B
membrane lipid strip
Triglyceride
Diacylglycerol
Phosphatidic Acid
Phosphatidylserine
Phosphatidylethanol-amine
Phosphatidylcholine
Phosphatidylglycerol
Cardiolipin
Phosphatidylinositol
PtdIns(4)P
PtdIns(4,5)P
PtdIns(3,4,5)P
Cholesterol
Sphingomyelin
3-sulfogalactosylcer-amide (Sulfatide)
Blank
BefA
AimA-His
BefA point mutants
R118E
R142E
K169A
R195A
W204A
Y232A
W251A
PS
CL
PIP lipid strip
Lysophosphatidic Acid
Lysophophocholine
PtdIns
PtdIns(3)P
PtdIns(4)P
PtdIns(5)P
Phophatidylethanol-amine
Phosphatidylcholine
Sphingosine-1-phophate
PtdIns(3,4)P2
PtdIns(3,5)P2
PtdIns(4,5)P2
PtdIns(3,4,5)P2
Phosphatidic Acid
Phosphatidylserine
Blank
BefA
Grip
AimA-His
PS
Pt-dIns(4,5)P
membrane lipid array
BefA 0 mM Ca
BefA 1 mM Ca
100 50 25 12.5 6.3 3.1 1.6
100 50 25 12.5 6.3 3.1 1.6
pmol/spot
Diacylglycerol
Phophatidic Acid
Phosphatidylserine
Phosphatidylinositol
Phosphatidylethanol-amine
Phosphatidylcholine
Phosphatidylglycerol
Sphingomyelin
C
concentration (feature(s)/nL)
0
1
2
3
4
5
control total
BefA total
control single vesicles
BefA single vesicles
control multi vesicles
BefA multi vesicles

### Figure S3

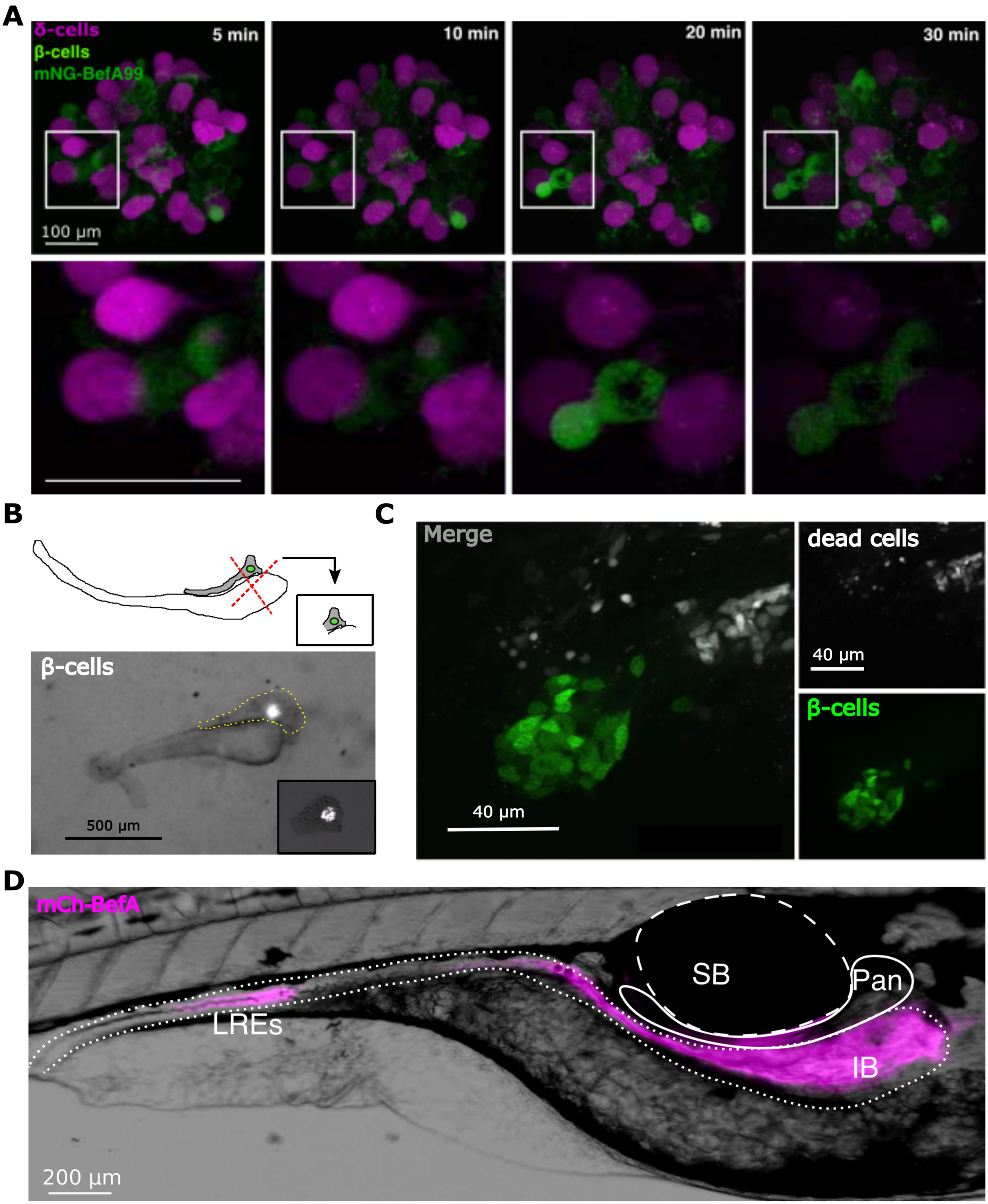
A
δ-cells
β-cells
mNG-BefA99
5 min
10 min
20 min
30 min
100 μm
B
β-cells
500 μm
C
Merge
40 μm
dead cells
40 μm
β-cells
D
mCh-BefA
SB
Pan
LREs
IB
200 μm

### Figure S4

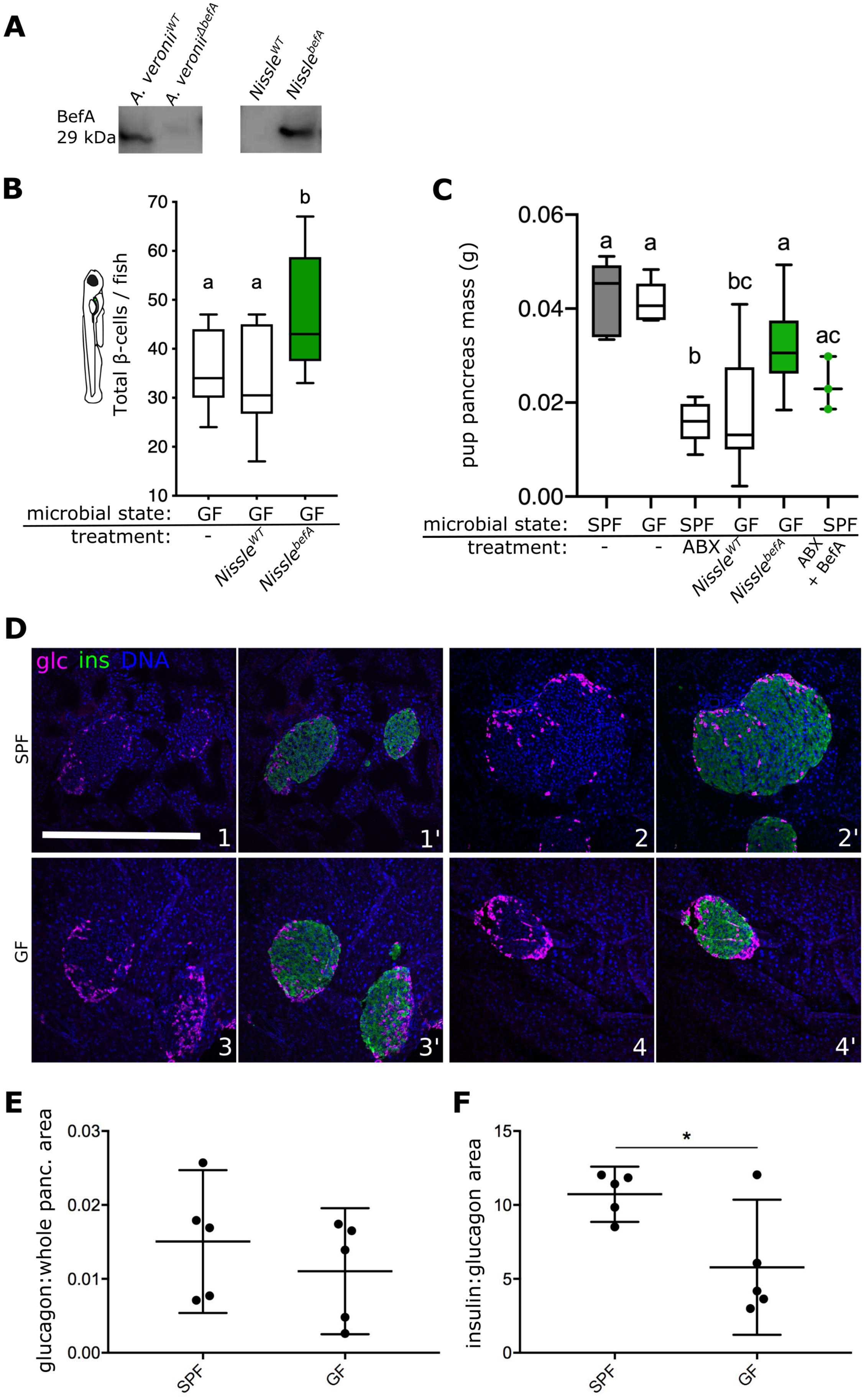

A
A. veronii$^{WT}$
A. veronii$^{\Delta befA}$
Nissle$^{WT}$
Nissle$^{befA}$
BefA
29 kDa
B
Total β-cells / fish
70
60
50
40
30
20
10
a
a
b
microbial state:
GF
GF
GF
treatment:
-
Nissle$^{WT}$
Nissle$^{befA}$
C
pup pancreas mass (g)
0.06
0.04
0.02
0.00
a
a
b
bc
a
ac
microbial state:
SPF
GF
SPF
GF
GF
SPF
treatment:
-
-
ABX
Nissle$^{WT}$
Nissle$^{befA}$
ABX + BefA
D
glc ins DNA
SPF
GF
1
1'
2
2'
3
3'
4
4'
E
glucagon:whole panc. area
0.03
0.02
0.01
0.00
SPF
GF
F
insulin:glucagon area
15
10
5
0
*
SPF
GF

#### Figure S5

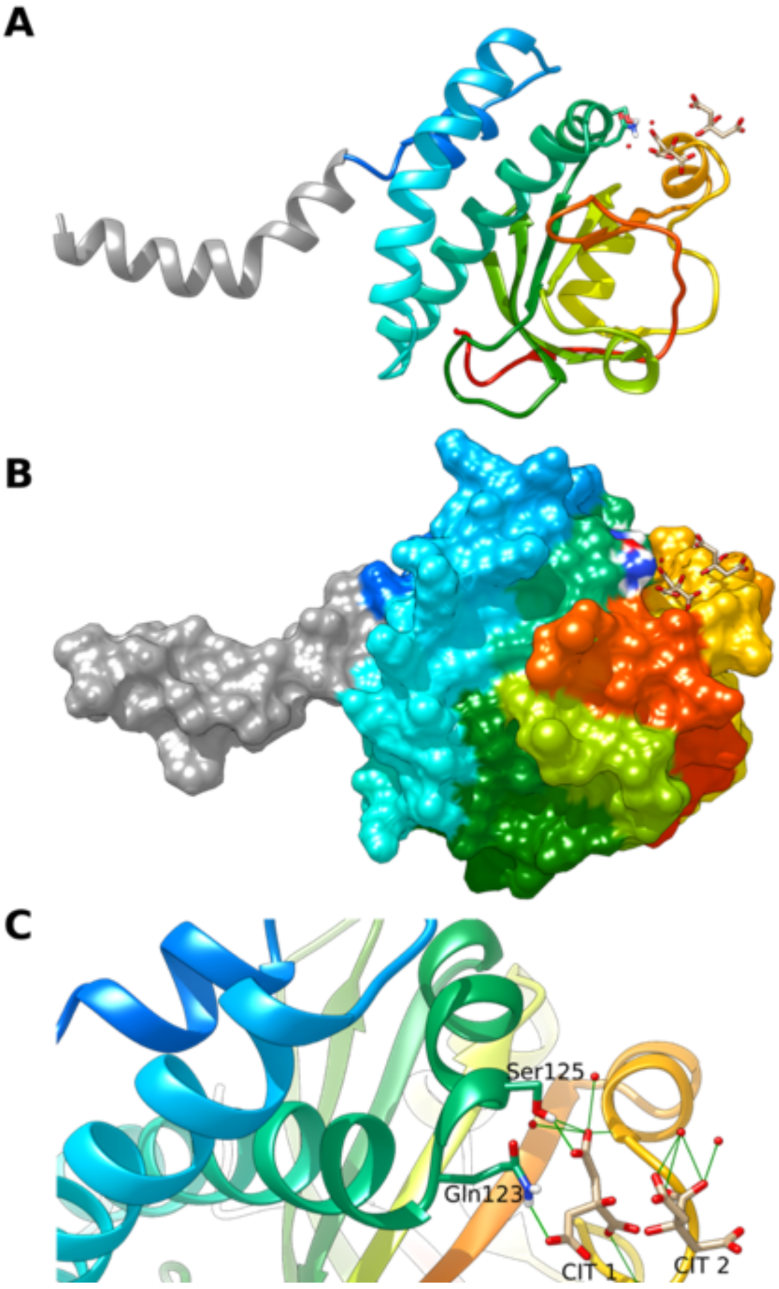

**Supplemental Figure Legends**

Figure S1. *Amino-acid alignments and phylogenetic analysis of diverse SYLF domain-containing proteins.*

(**A**) ClustalW alignment of amino acid sequences for the SYLF domains of four different SYLF domain-containing proteins: SH3YL1, YSC84, BefA, and the BefA homolog from *K. aerogenes*. The percentage values in the top left at the beginning of the sequences, indicate conserved amino acid identity to BefA across the sequence region shown. The corresponding secondary structure of BefA is shown across the top of the alignment, starting at H4 in teal. Pink triangles denote the predicted start site of the SYLF domain based on sequence alone. Purple boxes show residues involved in actin binding. Black underlines indicate partial conservation between BefA and key functional residues from YSC84. The numbers flanking the BefA SYLF sequence denote amino acid number. (**B&C**) 3D schematics of the bacterial BPSL1445 NMR structure (blue) (**B**) and the predicted structures for the SYLF domains of yeast protein YSC84 (gold), and human protein SH3YL1 (purple). Each one is overlayed with full length BefA structure (green). (**D**) A relatedness tree of SYLF homologs across bacterial genera. Homologs are grouped by genus and each branch represents the most closely related homolog to BefA within a genus. Bold taxa are annotated to a higher taxonomic rank than genus. Taxa are highlighted based on the environments where members within that clade have been found: green = genera found in aquatic hosts, grey = genera found in aquatic environments, yellow = genera found in terrestrial hosts, and pink = genera found in human hosts, the original BefA sequence from Aeromonas HM21 SYLF query and its close homolog from ZOR0001 are not highlighted, but instead have pink arrows for genera found in human hosts.

(**E**) The distribution of sequence lengths of putative SYLF-domain containing proteins is shifted in Eukaryotes compared to Prokaryotes. A relative frequency distribution (in percentages) of sequence lengths of proteins that contain an annotated SYLF-domain from Eukaryotes, Prokaryotes, and predicted BefA homologs.

*Figure S2. Exploration of the biochemical interactions of BefA with F-actin and membrane lipids*

(**A**) Coomasie Brilliant Blue denaturing protein gels illustrating results of F-actin co-pelleting assay. Ladder in lane 1 with band sizes in kDa labelled left. Expected band sizes for: BefA (29 kDa), $BefA^{99}$ (18 kDa), cortactin (80 kDa) and F-actin (42 kDa). + indicates the corresponding protein in left-hand label was added to the reaction shown in the lane immediately above. S = supernatant, P = pellet.

(**B**) Top row, Membrane lipid strip binding assay. From Left to right: legend, representative strip incubated with BefA shows positive binding to phosphatidylserine (PS) and cardiolipin (CL), negative control performed with immunomodulatory bacterial protein His-AimA, point mutants created within individual predicted lipid binding sites (amino acid change indicated by label). Middle row, PIP lipid strip binding assay. From left to right: legend, representative strip incubated with BefA shows positive binding to PS, positive control was performed with $PI(4,5)P_2$ Grip and bound to $PI(4,5)P_2$ as expected (*denotes positive control of HRP secondary antibody spotted directly onto strip above Sphingosine-1-phosphate to confirm visualizing reagents were working), negative control with His-AimA. Bottom row, Membrane lipid array assay was performed with BefA with or without 1 mM Ca.

(**C**) Concentration of total fluorescent liposomes with separate quantifications of total single and multi-vesicle structures, illustrating untreated groups do not form multi-vesicle structures.

Figure S3. *Tracking the activity of BefA in zebrafish*

(**A**) Fluorescent images of a single primary zebrafish islet treated with mNG-BefA$^{99}$ over the course of 30 minutes. magenta = δ-cells, green = mNG and β-cells. Top row from left to right: whole islet images 5-,10-, 15-, 20- and 30-minutes post addition of mNG-BefA$^{99}$. Scale bar = 100 μm. boxes outline inset of image at higher magnification in bottom row directly below. Scale bar = 100 μm.

(**B**) Top: cartoon representation of a dissected larval intestine (white) with attached pancreas (grey) and primary islet indicated in green. Red dashed lines illustrate the approximate dissections to separate the head of the pancreas from the rest of the tissue. Bottom: greyscale image of an actual dissection. Pancreas (outlined in dashed yellow line) attached above the intestine, with the final resulting pancreatic head containing in-tact islet in lower right inset. Insulin promoter expressing β-cells in white. Scale bar = 500 μm.

(**C**) Confocal image of β-cells in explant tissue after 48 hrs of ex-vivo culture. TOPRO staining denoting dead or dying cells in white (top left), insulin promoter expressing β-cells in green (bottom left). Merge = right. Scale bars = 40 μm.

(**D**) DIC image of 4 dpf larval zebrafish following oral microgavage treatment with mCh-BefA. LREs = Lysozyme Rich Enterocytes of the distal intestine. For anatomical orientation: IB = intestinal bulb, SB = swim bladder, Pan = pancreas. Scale bar = 200 µm.

*Figure S4. Activity of BefA is conserved in mice*

(**A**) Western blots of anti-BefA antibody binding to CFS from indicated bacterial cultures.

(**B**) Boxplots representing total beta-cell per GF larvae monoassociated with Nissle strains.

(**C**) Boxplots illustrating total pancreas mass in grams (g) for neonatal P12 aged mice used in Figure 4A.

(**D**) Representative images of immunofluorescence stained pancreatic tissue from four different Swiss Webster mice (1-4). Top row: SPF mice, Bottom row: GF mice. Magenta = Glucagon (glc) denoting alpha cells. Green = insulin (ins) denoting beta-cells (only in1'-4'), blue = DAPI denoting nuclei. Scale bar = 500 µm.

(**E**) Average ratio of alpha-cell area to whole pancreas area per mouse.

(**F**) Average ratio of alpha-cell area to beta-cell area per mouse.

**Table S1.**

Table S1. Data Collection and Refinement Statistics for Deposited Model of BefA

| **Data Collection** | **Selenomet** | **native** |
|---|---|---|
| space group | C121 | C121 |
| unit cell a, b, c (Å) | 94.5, 64.3,42.6 | 94.7, 64.5, 42.6 |
| alpha, beta, gamma (degrees) | 90, 113.4, 90 | 90, 113.3, 90 |
| resolution (Å) | 50-1.29 (1.31-1.29) | 37.41-1.27 (1.34-1.27) |
| completeness (%) | 96.6 (92.7) | 100 (99.8) |

| | | |
|---|---|---|
| no. unique reflections | 56577 | 62064 (6164) |
| multiplicity | 6.1 (4.0) | 10 (6.9) |
| <I/sigma> | 24.7 (1.2) | 8.7 (3.2) |
| CC1/2 (outer shell) | 0.45 | 0.994 (0.341) |
| R merge | 0.097 (1.25) | 0.168 (1.42) |
| **Refinement** | | |
| R work (%) | | 13.2 |
| R free (%) | | 15.4 |
| no. of molecules in the asymmetric unit | | 1 |
| no. protein residues | | 223 |
| no. of waters | | 216 |
| rmsd for lengths (Å) | | 0.014 |
| rmsd for angles (deg) | | 1.46 |
| Ramachandran plot (%) | | |
| preferred | | 97.5 |
| allowed | | 1.3 |
| outliers | | 1.2 |
| Avg. B factor (Å^2) | | |
| mainchain[A] | | 13 |
| waters | | 33 |
| new PDB entry | | 7RFQ |

### Full wwPDB X-ray Structure Validation Report (i)

Jul 15, 2021 – 10:38 AM EDT

| | | |
|---|---|---|
| PDB ID | : | 7RFQ |
| Title | : | STRUCTURE OF BACTERIAL SYLF DOMAIN CONTAINING PROTEIN, BETA CELL EXPANSION FACTOR A (BEFA) |
| Deposited on | : | 2021-07-14 |
| Resolution | : | 1.27 Å(reported) |

This is a Full wwPDB X-ray Structure Validation Report.

This report is produced by the wwPDB biocuration pipeline after annotation of the structure.

We welcome your comments at **
A user guide is available at
https://www.wwpdb.org/validation/2017/XrayValidationReportHelp
with specific help available everywhere you see the (i) symbol.

The following versions of software and data (see references (i)) were used in the production of this report:

| | | |
|---|---|---|
| MolProbity | : | 4.02b-467 |
| Mogul | : | 1.8.5 (274361), CSD as541be (2020) |
| Xtriage (Phenix) | : | 1.13 |
| EDS | : | 2.22 |
| Percentile statistics | : | 20191225.v01 (using entries in the PDB archive December 25th 2019) |
| Refmac | : | 5.8.0158 |
| CCP4 | : | 7.0.044 (Gargrove) |
| Ideal geometry (proteins) | : | Engh & Huber (2001) |
| Ideal geometry (DNA, RNA) | : | Parkinson et al. (1996) |
| Validation Pipeline (wwPDB-VP) | : | 2.22 |

### 1 Overall quality at a glance (i)

The following experimental techniques were used to determine the structure:
*X-RAY DIFFRACTION*

The reported resolution of this entry is 1.27 Å.

Percentile scores (ranging between 0-100) for global validation metrics of the entry are shown in the following graphic. The table shows the number of entries on which the scores are based.

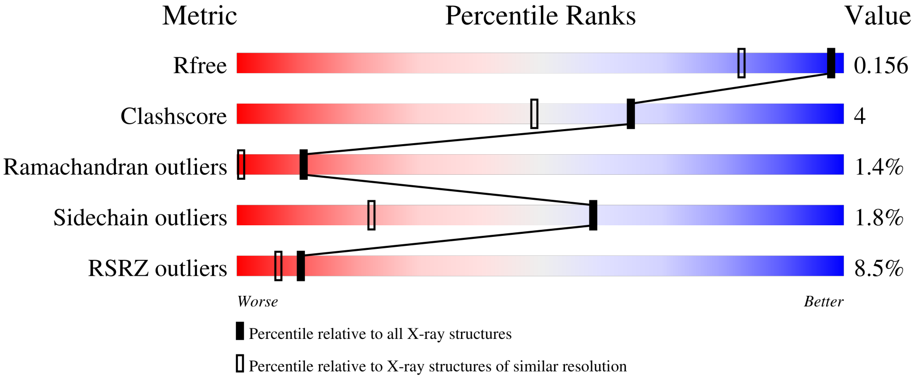


| Metric | Whole archive (#Entries) | Similar resolution (#Entries, resolution range(Å)) |
|---|---|---|
| $R_{free}$ | 130704 | 1850 (1.30-1.26) |
| Clashscore | 141614 | 1926 (1.30-1.26) |
| Ramachandran outliers | 138981 | 1860 (1.30-1.26) |
| Sidechain outliers | 138945 | 1859 (1.30-1.26) |
| RSRZ outliers | 127900 | 1807 (1.30-1.26) |

The table below summarises the geometric issues observed across the polymeric chains and their fit to the electron density. The red, orange, yellow and green segments of the lower bar indicate the fraction of residues that contain outliers for >=3, 2, 1 and 0 types of geometric quality criteria respectively. A grey segment represents the fraction of residues that are not modelled. The numeric value for each fraction is indicated below the corresponding segment, with a dot representing fractions <=5% The upper red bar (where present) indicates the fraction of residues that have poor fit to the electron density. The numeric value is given above the bar.

| Mol | Chain | Length | Quality of chain |
|---|---|---|---|
| 1 | A | 270 | 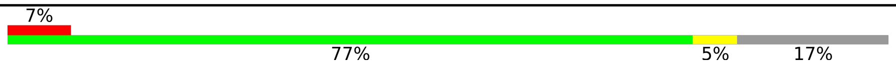 |

### 2 Entry composition (i)

There are 3 unique types of molecules in this entry. The entry contains 3819 atoms, of which 1751 are hydrogens and 0 are deuteriums.

In the tables below, the ZeroOcc column contains the number of atoms modelled with zero occupancy, the AltConf column contains the number of residues with at least one atom in alternate conformation and the Trace column contains the number of residues modelled with at most 2 atoms.

- Molecule 1 is a protein called BETA CELL EXPANSION FACTOR A (BEFA).

| Mol | Chain | Residues | Atoms | | | | | | ZeroOcc | AltConf | Trace |
|---|---|---|---|---|---|---|---|---|---|---|---|
| | | | Total | C | H | N | O | S | | | |
| 1 | A | 223 | 3561 | 1163 | 1741 | 296 | 352 | 9 | 0 | 25 | 0 |

There are 9 discrepancies between the modelled and reference sequences:

| Chain | Residue | Modelled | Actual | Comment | Reference |
|---|---|---|---|---|---|
| A | 0 | MET | - | initiating methionine | UNP A0A2S3XLU2 |
| A | 262 | LEU | - | expression tag | UNP A0A2S3XLU2 |
| A | 263 | GLU | - | expression tag | UNP A0A2S3XLU2 |
| A | 264 | HIS | - | expression tag | UNP A0A2S3XLU2 |
| A | 265 | HIS | - | expression tag | UNP A0A2S3XLU2 |
| A | 266 | HIS | - | expression tag | UNP A0A2S3XLU2 |
| A | 267 | HIS | - | expression tag | UNP A0A2S3XLU2 |
| A | 268 | HIS | - | expression tag | UNP A0A2S3XLU2 |
| A | 269 | HIS | - | expression tag | UNP A0A2S3XLU2 |

- Molecule 2 is CITRIC ACID (three-letter code: CIT) (formula: $C_6H_8O_7$).

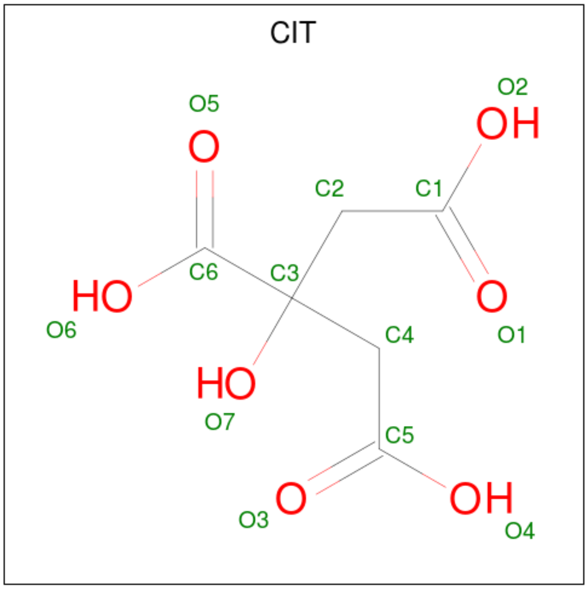

| Mol | Chain | Residues | Atoms | ZeroOcc | AltConf |
|---|---|---|---|---|---|
| 2 | A | 1 | Total C H O<br>18 6 5 7 | 0 | 0 |
| 2 | A | 1 | Total C H O<br>18 6 5 7 | 0 | 0 |

- Molecule 3 is water.

| Mol | Chain | Residues | Atoms | ZeroOcc | AltConf |
|---|---|---|---|---|---|
| 3 | A | 213 | Total O<br>222 222 | 0 | 9 |

### 3 Residue-property plots ⓘ

These plots are drawn for all protein, RNA, DNA and oligosaccharide chains in the entry. The first graphic for a chain summarises the proportions of the various outlier classes displayed in the second graphic. The second graphic shows the sequence view annotated by issues in geometry and electron density. Residues are color-coded according to the number of geometric quality criteria for which they contain at least one outlier: green = 0, yellow = 1, orange = 2 and red = 3 or more. A red dot above a residue indicates a poor fit to the electron density (RSRZ > 2). Stretches of 2 or more consecutive residues without any outlier are shown as a green connector. Residues present in the sample, but not in the model, are shown in grey.

• Molecule 1: BETA CELL EXPANSION FACTOR A (BEFA)

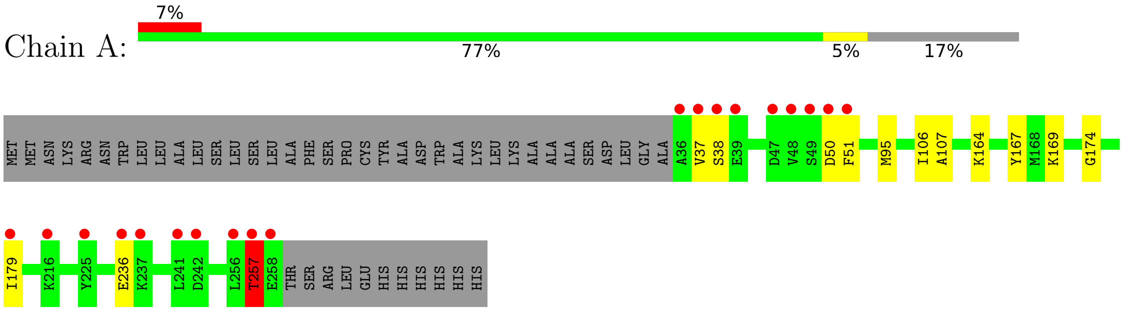

### 4 Data and refinement statistics (i)

| Property | Value | Source |
|---|---|---|
| Space group | C 1 2 1 | Depositor |
| Cell constants<br>a, b, c, $\alpha$, $\beta$, $\gamma$ | 94.67Å 64.55Å 42.63Å<br>90.00° 113.26° 90.00° | Depositor |
| Resolution (Å) | 37.35 – 1.27<br>37.35 – 1.27 | Depositor<br>EDS |
| % Data completeness<br>(in resolution range) | 99.9 (37.35-1.27)<br>99.9 (37.35-1.27) | Depositor<br>EDS |
| $R_{merge}$ | (Not available) | Depositor |
| $R_{sym}$ | 0.17 | Depositor |
| $< I/\sigma(I) >$ [1] | 2.47 (at 1.27Å) | Xtriage |
| Refinement program | PHENIX 1.9_1692 | Depositor |
| R, $R_{free}$ | 0.131 , 0.155<br>0.132 , 0.156 | Depositor<br>DCC |
| $R_{free}$ test set | 2308 reflections (3.72%) | wwPDB-VP |
| Wilson B-factor (Å$^2$) | 10.8 | Xtriage |
| Anisotropy | 0.181 | Xtriage |
| Bulk solvent $k_{sol}$(e/Å$^3$), $B_{sol}$(Å$^2$) | 0.42 , 58.6 | EDS |
| L-test for twinning[2] | $< \|L\| >$ = 0.48, $< L^2 >$ = 0.31 | Xtriage |
| Estimated twinning fraction | No twinning to report. | Xtriage |
| $F_o$,$F_c$ correlation | 0.98 | EDS |
| Total number of atoms | 3819 | wwPDB-VP |
| Average B, all atoms (Å$^2$) | 18.0 | wwPDB-VP |

Xtriage's analysis on translational NCS is as follows: *The largest off-origin peak in the Patterson function is 9.14% of the height of the origin peak. No significant pseudotranslation is detected.*

[1]Intensities estimated from amplitudes.

[2]Theoretical values of $< |L| >$, $< L^2 >$ for acentric reflections are 0.5, 0.333 respectively for untwinned datasets, and 0.375, 0.2 for perfectly twinned datasets.

### 5 Model quality (i)

#### 5.1 Standard geometry (i)

Bond lengths and bond angles in the following residue types are not validated in this section: CIT

The Z score for a bond length (or angle) is the number of standard deviations the observed value is removed from the expected value. A bond length (or angle) with $|Z| > 5$ is considered an outlier worth inspection. RMSZ is the root-mean-square of all Z scores of the bond lengths (or angles).

| Mol | Chain | Bond lengths | | Bond angles | |
|---|---|---|---|---|---|
| | | RMSZ | #\|Z\| >5 | RMSZ | #\|Z\| >5 |
| 1 | A | 0.64 | 0/1913 | 0.67 | 0/2586 |

There are no bond length outliers.

There are no bond angle outliers.

There are no chirality outliers.

There are no planarity outliers.

#### 5.2 Too-close contacts (i)

In the following table, the Non-H and H(model) columns list the number of non-hydrogen atoms and hydrogen atoms in the chain respectively. The H(added) column lists the number of hydrogen atoms added and optimized by MolProbity. The Clashes column lists the number of clashes within the asymmetric unit, whereas Symm-Clashes lists symmetry-related clashes.

| Mol | Chain | Non-H | H(model) | H(added) | Clashes | Symm-Clashes |
|---|---|---|---|---|---|---|
| 1 | A | 1820 | 1741 | 1735 | 14 | 0 |
| 2 | A | 26 | 10 | 10 | 1 | 0 |
| 3 | A | 222 | 0 | 0 | 2 | 0 |
| All | All | 2068 | 1751 | 1745 | 15 | 0 |

The all-atom clashscore is defined as the number of clashes found per 1000 atoms (including hydrogen atoms). The all-atom clashscore for this structure is 4.

All (15) close contacts within the same asymmetric unit are listed below, sorted by their clash magnitude.

| Atom-1 | Atom-2 | Interatomic distance (Å) | Clash overlap (Å) |
|---|---|---|---|
| 1:A:95[B]:MET:HE1 | 1:A:106:ILE:HG22 | 1.78 | 0.65 |

*Continued on next page...*

*Continued from previous page...*

| Atom-1 | Atom-2 | Interatomic distance (Å) | Clash overlap (Å) |
|---|---|---|---|
| 1:A:169[B]:LYS:HG2 | 3:A:519[B]:HOH:O | 2.11 | 0.50 |
| 1:A:167:TYR:H | 1:A:257[A]:THR:HG23 | 1.79 | 0.48 |
| 1:A:95[B]:MET:CE | 1:A:106:ILE:HG22 | 2.44 | 0.48 |
| 1:A:95[B]:MET:CE | 1:A:107:ALA:HA | 2.47 | 0.45 |
| 1:A:174:GLY:O | 1:A:179:ILE:HG13 | 2.18 | 0.43 |
| 2:A:301:CIT:O6 | 2:A:302:CIT:O5 | 2.38 | 0.41 |
| 1:A:164:LYS:NZ | 3:A:413:HOH:O | 2.54 | 0.41 |

There are no symmetry-related clashes.

#### 5.3 Torsion angles (i)

##### 5.3.1 Protein backbone (i)

In the following table, the Percentiles column shows the percent Ramachandran outliers of the chain as a percentile score with respect to all X-ray entries followed by that with respect to entries of similar resolution.

The Analysed column shows the number of residues for which the backbone conformation was analysed, and the total number of residues.

| Mol | Chain | Analysed | Favoured | Allowed | Outliers | Percentiles |
|---|---|---|---|---|---|---|
| 1 | A | 246/270 (91%) | 235 (96%) | 5 (2%) | 6 (2%) | 6 0 |

All (6) Ramachandran outliers are listed below:

| Mol | Chain | Res | Type |
|---|---|---|---|
| 1 | A | 51[A] | PHE |
| 1 | A | 51[B] | PHE |
| 1 | A | 236[A] | GLU |
| 1 | A | 236[B] | GLU |
| 1 | A | 257[A] | THR |
| 1 | A | 257[B] | THR |

##### 5.3.2 Protein sidechains (i)

In the following table, the Percentiles column shows the percent sidechain outliers of the chain as a percentile score with respect to all X-ray entries followed by that with respect to entries of similar resolution.

The Analysed column shows the number of residues for which the sidechain conformation was analysed, and the total number of residues.

| Mol | Chain | Analysed | Rotameric | Outliers | Percentiles | |
|---|---|---|---|---|---|---|
| 1 | A | 186/212 (88%) | 182 (98%) | 4 (2%) | 52 | 14 |

All (4) residues with a non-rotameric sidechain are listed below:

| Mol | Chain | Res | Type |
|---|---|---|---|
| 1 | A | 37 | VAL |
| 1 | A | 38 | SER |
| 1 | A | 257[A] | THR |
| 1 | A | 257[B] | THR |

Sometimes sidechains can be flipped to improve hydrogen bonding and reduce clashes. There are no such sidechains identified.

##### 5.3.3 RNA (i)

There are no RNA molecules in this entry.

#### 5.4 Non-standard residues in protein, DNA, RNA chains (i)

There are no non-standard protein/DNA/RNA residues in this entry.

#### 5.5 Carbohydrates (i)

There are no monosaccharides in this entry.

#### 5.6 Ligand geometry (i)

2 ligands are modelled in this entry.

In the following table, the Counts columns list the number of bonds (or angles) for which Mogul statistics could be retrieved, the number of bonds (or angles) that are observed in the model and the number of bonds (or angles) that are defined in the Chemical Component Dictionary. The Link column lists molecule types, if any, to which the group is linked. The Z score for a bond length (or angle) is the number of standard deviations the observed value is removed from the expected value. A bond length (or angle) with $|Z| > 2$ is considered an outlier worth inspection. RMSZ is the root-mean-square of all Z scores of the bond lengths (or angles).

| Mol | Type | Chain | Res | Link | Bond lengths | | | Bond angles | | |
|---|---|---|---|---|---|---|---|---|---|---|
| | | | | | Counts | RMSZ | #\|Z\| > 2 | Counts | RMSZ | #\|Z\| > 2 |
| 2 | CIT | A | 301 | - | 3,12,12 | 1.62 | 1 (33%) | 3,17,17 | 1.43 | 1 (33%) |
| 2 | CIT | A | 302 | - | 3,12,12 | 1.05 | 0 | 3,17,17 | 1.96 | 1 (33%) |

In the following table, the Chirals column lists the number of chiral outliers, the number of chiral centers analysed, the number of these observed in the model and the number defined in the Chemical Component Dictionary. Similar counts are reported in the Torsion and Rings columns. '-' means no outliers of that kind were identified.

| Mol | Type | Chain | Res | Link | Chirals | Torsions | Rings |
|---|---|---|---|---|---|---|---|
| 2 | CIT | A | 301 | - | - | 0/6/16/16 | - |
| 2 | CIT | A | 302 | - | - | 1/6/16/16 | - |

All (1) bond length outliers are listed below:

| Mol | Chain | Res | Type | Atoms | Z | Observed(Å) | Ideal(Å) |
|---|---|---|---|---|---|---|---|
| 2 | A | 301 | CIT | C2-C3 | -2.45 | 1.51 | 1.54 |

All (2) bond angle outliers are listed below:

| Mol | Chain | Res | Type | Atoms | Z | Observed($^o$) | Ideal($^o$) |
|---|---|---|---|---|---|---|---|
| 2 | A | 302 | CIT | C3-C2-C1 | -3.27 | 109.75 | 114.98 |
| 2 | A | 301 | CIT | C3-C2-C1 | 2.04 | 118.26 | 114.98 |

There are no chirality outliers.

All (1) torsion outliers are listed below:

| Mol | Chain | Res | Type | Atoms |
|---|---|---|---|---|
| 2 | A | 302 | CIT | C1-C2-C3-O7 |

There are no ring outliers.

2 monomers are involved in 1 short contact:

| Mol | Chain | Res | Type | Clashes | Symm-Clashes |
|---|---|---|---|---|---|
| 2 | A | 301 | CIT | 1 | 0 |
| 2 | A | 302 | CIT | 1 | 0 |

#### 5.7 Other polymers (i)

There are no such residues in this entry.

#### 5.8 Polymer linkage issues (i)

There are no chain breaks in this entry.

### 6 Fit of model and data (i)

#### 6.1 Protein, DNA and RNA chains (i)

In the following table, the column labelled '#RSRZ> 2' contains the number (and percentage) of RSRZ outliers, followed by percent RSRZ outliers for the chain as percentile scores relative to all X-ray entries and entries of similar resolution. The OWAB column contains the minimum, median, $95^{th}$ percentile and maximum values of the occupancy-weighted average B-factor per residue. The column labelled 'Q< 0.9' lists the number of (and percentage) of residues with an average occupancy less than 0.9.

| Mol | Chain | Analysed | <RSRZ> | #RSRZ>2 | | | OWAB(Å$^2$) | Q<0.9 |
|---|---|---|---|---|---|---|---|---|
| 1 | A | 223/270 (82%) | 0.54 | 19 (8%) | 10 | 7 | 7, 14, 29, 47 | 0 |

All (19) RSRZ outliers are listed below:

| Mol | Chain | Res | Type | RSRZ |
|---|---|---|---|---|
| 1 | A | 37 | VAL | 8.7 |
| 1 | A | 51[A] | PHE | 7.0 |
| 1 | A | 36 | ALA | 6.8 |
| 1 | A | 258 | GLU | 6.4 |
| 1 | A | 257[A] | THR | 5.1 |
| 1 | A | 237 | LYS | 5.0 |
| 1 | A | 38 | SER | 4.8 |
| 1 | A | 50[A] | ASP | 4.5 |
| 1 | A | 47[A] | ASP | 3.8 |
| 1 | A | 48[A] | VAL | 3.6 |
| 1 | A | 225 | TYR | 3.4 |
| 1 | A | 242 | ASP | 3.3 |
| 1 | A | 49 | SER | 3.1 |
| 1 | A | 241 | LEU | 3.0 |
| 1 | A | 179 | ILE | 2.9 |
| 1 | A | 39 | GLU | 2.8 |
| 1 | A | 236[A] | GLU | 2.7 |
| 1 | A | 256[A] | LEU | 2.6 |
| 1 | A | 216 | LYS | 2.2 |

#### 6.2 Non-standard residues in protein, DNA, RNA chains (i)

There are no non-standard protein/DNA/RNA residues in this entry.

#### 6.3 Carbohydrates (i)

There are no monosaccharides in this entry.

#### 6.4 Ligands (i)

In the following table, the Atoms column lists the number of modelled atoms in the group and the number defined in the chemical component dictionary. The B-factors column lists the minimum, median, $95^{th}$ percentile and maximum values of B factors of atoms in the group. The column labelled 'Q< 0.9' lists the number of atoms with occupancy less than 0.9.

| Mol | Type | Chain | Res | Atoms | RSCC | RSR | B-factors(Å$^2$) | Q<0.9 |
|---|---|---|---|---|---|---|---|---|
| 2 | CIT | A | 302 | 13/13 | 0.70 | 0.23 | 38,44,55,55 | 0 |
| 2 | CIT | A | 301 | 13/13 | 0.86 | 0.19 | 30,34,48,48 | 0 |

#### 6.5 Other polymers (i)

There are no such residues in this entry.
